# Integrative single-cell analysis of neural stem/progenitor cells reveals epigenetically dysregulated interferon response in progressive multiple sclerosis

**DOI:** 10.1101/2024.02.09.579648

**Authors:** Bongsoo Park, Alexandra M. Nicaise, Dimitrios Tsitsipatis, Liviu Pirvan, Pranathi Prasad, Miguel Larraz Lopez De Novales, Julia Whitten, Luka Culig, Joseph Llewellyn, Rosana-Bristena Ionescu, Cory M. Willis, Grzegorz Krzak, Jinshui Fan, Supriyo De, Marta Suarez Cubero, Angeliki Spathopoulou, Luca Peruzzotti-Jametti, Tommaso Leonardi, Frank Edenhofer, Myriam Gorospe, Irina Mohorianu, Stefano Pluchino, Isabel Beerman

## Abstract

Progressive multiple sclerosis (PMS) is characterized by a primary smouldering pathological disease process associated with a superimposed inflammatory activity. Cellular and molecular processes sustaining the pathobiology of PMS remain to be identified.

We previously discovered senescence signatures in neural stem/progenitor cells (NSCs) from people with PMS. Applying direct reprogramming to generate directly induced NSCs (iNSCs) from somatic fibroblasts, we retain epigenetic information and observe hypomethylation of genes associated with lipid metabolic processes and IFN signalling only in PMS lines. Single-cell/nucleus transcriptomic and epigenetic profiling reveal an inflammatory, senescent-like, IFN-responsive radial glia (RG)-like cell subcluster mainly in PMS iNSCs that is driven by IFN-associated transcription factors. Lastly, we identify a population of senescent, IFN-responsive, disease-associated RG-like cells (DARGs) in the PMS brain that share pseudotime trajectories with iNSCs *in vitro*.

We describe the existence of a non-neurogenic, dysfunctional DARG population that has the potential to fuel smouldering inflammation in PMS.

## INTRODUCTION

Multiple sclerosis (MS) is a complex neuroinflammatory and neurodegenerative disease characterized by inflammation and demyelination in the central nervous system (CNS). It is believed to be caused by an interplay of genetic predisposition and environmental risk factors, such as exposure to viruses.^1^ The early phase of the disease, known as relapsing remitting MS (RRMS), pathologically manifests as acute demyelinating lesions with some endogenous repair. There are disease-modifying therapies (DMTs) available that target peripheral immune cells to reduce the risk of developing new lesions and clinical relapses. Over time, however, most people with RRMS transition into a progressive (PMS) stage of the disease that is characterized by the steady accumulation of neurological disability in the absence of endogenous repair that leads to neurodegeneration. Despite most available DMTs being effective in people with RRMS they are much less effective in people with PMS. This has made the progressive stage of the disease an unmet clinical need.^1^

Pathological, neuroimaging, and clinical data suggest that progressive MS is driven by a primary smouldering process associated with inflammation. Several mechanisms have been proposed to drive smouldering MS, including innate immune activation, demyelination and energy deficits, adaptive immunity, and, recently, age-related mechanisms.^2,3^

Age is one of the most significant risk factors in the development of PMS, which is similar to other neurodegenerative diseases such as Alzheimer’s and Parkinson’s disease.^4,5^ Longitudinal assessment of brain aging using imaging technologies showed that people with MS demonstrate increased ‘*brain age*’ compared to healthy controls that was defined by increased atrophy and decreased grey matter volumes.^6,7^ Pathological hallmarks of cellular aging, such as senescence and senescence-associated changes, have been identified in people with PMS. These include decreased telomere length in peripheral leukocytes,^8–10^ increased DNA and mitochondrial damage in neurons *in situ,*^11–13^ increased epigenetic age in glial cells,^14^ senescence-associated secretory phenotype (SASP) in microglia and astrocytes,^15,16^ and p16/CDKN2A expression in glia and neural stem/progenitor cells (NSCs).^16,17^ The increasing body of evidence suggesting an association of PMS with cellular senescence requires further study to understand how the accumulation of senescent glial cells contributes to disease pathogenesis.

The genetic components underlying MS risk and severity are incredibly complex. Genome wide association studies (GWAS) largely implicated cells of the peripheral immune system in the development of disease. However, mapping of MS susceptibility genes onto brain tissue has revealed enrichment of MS-susceptibility genes in glial cells of the CNS, including astrocytes, oligodendrocytes, and microglia. Moreover, prediction of MS severity from genetic loci involved in the CNS were found to be associated with mitochondrial function, cellular senescence, and synaptic plasticity.^18^

Recent work with induced pluripotent stem cells (iPSCs), which retain the genetic information from the starting cell and is maintained in the resulting cell type of interest, has allowed for modelling of complex human brain disorders *in vitro*. Although iPSCs offer a powerful platform for modelling of human diseases, epigenetic modifications, especially those associated with aging, are lost in the reprogramming process due to the use of the Yamanaka factors.^19–21^ Epigenetic changes, which can be reflected by environmental risk factors such as smoking and exposure to viruses, are likely contribute to MS susceptibility and progression. A few studies have identified aberrant DNA methylation patterns in both CNS tissue and peripheral blood and leukocytes in people with MS. Many of these patterns are associated with functional pathways related to immune response, neuronal survival^22^, and demyelination.^23^ Recent work has found that iPSC-NSCs from patients with PMS display markers of senescence and prevent oligodendrocyte progenitor cell differentiation via the SASP.^17,24^ Furthermore, iPSC-astrocytes from MS patients also display senescence-related gene expression, dysfunctional metabolism, and increased immune and inflammatory genes.^25,26^

Stem cell exhaustion is a known hallmark of biological aging that is typically associated with reduced tissue repair. Adult brain stem cells/NSCs, also known as radial glia-like cells (RG), are astroglial-like cells that classically reside in the mammalian subventricular zone and dentate gyrus of the hippocampus and can give rise to mature neurons, astrocytes, and oligodendrocytes.^27^ Furthermore, noncanonical niches for RG have also been reported, including the neocortex of primates,^28^ the cerebellum of rabbits,^29^ the amygdala of mice,^30^ and the striatum of humans.^31^ Studies in animals have shown that with progressing age and neurodegenerative disease the capabilities of RGs, such as differentiation into neurons and repair capabilities.^32–34^ Many studies have attempted to address the existence, persistence, and role of NSC/RGs in the human brain following early post-natal life. Immunostaining of post-mortem human brains^35^ identified NSCs in the lining of the walls of the lateral ventricle, the dentate gyrus, and the olfactory epithelium.^36,37^ However, their capacity for neurogenesis and differentiation, as well as their presence outside of the main CNS germinal stem cell niches is much debated.^38,39^ More recently, phenomena such as inflammation and injury support both de-maturation of neurons^40^ and de-differentiation of astrocytes^41^ into NSC/RG-like cell states. NSCs expressing glial markers and displaying features of senescence have also been identified near lesioned areas in post-mortem MS brain tissue.^17,42,43^ Their role and putative function in brain physiology and disease is not currently understood.

Disease modelling with stem cell technologies holds the promise for understanding cellular dynamics that were previously unreachable in brain cells *in vivo*. We employed the use of direct reprogramming technology, which better retains epigenetic memory of the donor cells,^44^ to better investigate the origin of the senescent phenotype in NSCs within the context of PMS.^17,24,25^, We directly converted skin-derived fibroblasts from healthy human controls and people with PMS into stably expandable induced NSCs (iNSC). This was done by exposing fibroblasts to a transient (24-hour long) exposure to the Yamanaka factors^45^ in the presence of NSC differentiation factors to generate a heterogenous population of stem and progenitor cells.

By subjecting both the parental fibroblasts and iNSCs to whole genome bisulfite sequencing (WGBS), we identified hypomethylated genes encoding proteins that function in pathways similar to those associated with inflammatory and interferon (IFN) signalling in PMS cells, suggesting a predisposition to inflammation. Furthermore, direct reprogramming to iNSCs maintained epigenetic information from the donor cells. Through both bulk and single-cell/ nucleus transcriptomics analyses, we found increased activation of pathways pertaining to cellular senescence, inflammation, and IFN signalling in a subset of PMS iNSCs. Combined single-nuclei ATACseq supported changes in chromatin accessibility and was linked to consistent pathways. Within the heterogenous iNSCs, we identified a focused inflammatory and senescent-like cluster and elevated expression of related upstream transcription factor IRF1 in PMS iNSCs. We integrated published post-mortem single-cell/nuclei transcriptomics data sets and confirmed the presence of non-neurogenic, disease-associated RG-like cells (DARGs) in the PMS brain, primarily in chronic active, slowly expanding lesions, exhibiting senescence and IFN-responsive characteristics.

Our work confirms that direct reprogramming technology is a powerful tool to model disease-in-a-dish to study neurodegenerative disorders. In doing so, it led us to identify the existence of a long-neglected, non-neurogenic DARG cell cluster especially in chronic brain MS lesions that has the potential to fuel continuous smouldering inflammation in PMS.

## RESULTS

### Bulk (multi-modal) sequencing reveals upregulation of senescence and inflammatory pathways in PMS iNSCs

To model disease and age associated NSC features, we directly reprogrammed skin-derived fibroblasts from control healthy subjects (Ctrl) and people with PMS (PMS) into stably expandable iNSCs (**Table S1**).^45^ We confirmed the expression of established and accepted NSC markers, including the mRNAs, using RT-PCR, and proteins, using immunocytochemistry, that encode Nestin (NES), SOX2, ETNPPL, and PAX6, as well as the clearance of Sendai virus markers.^46^ PCA summary of transcriptomic signatures across samples (**Fig. 1A**) revealed robust signal across replicates and a clear separation between the Ctrl and PMS samples. Additional checks comprised incremental Jaccard similarity index (**Fig. S1A**),^47^ and assessment and removal of technical noise using noisyR (**Fig. S1B**),^48^ followed by normalization of expression levels.^47^ The differential expression analyses (with convergent results on the DESeq2^49^ and edgeR^50^ pipelines) led to 1,021 upregulated and 844 downregulated genes (FDR < 0.05, |log2(FC)| ≥ 0.5) that included transcripts related to senescence, such as *CDKN1A*, *IRF7*, and *ISG15* (**Fig. 1B**). Gene set enrichment analysis (GSEA^51^), using genes expressed above noise level as background, identified several pathways enriched in the PMS iNSCs, including mRNAs encoded by gene sets that drive response to stress (*TNC*, *STAT6*), immune system processes (*THBS1*, *SPP1*), positive regulation of lipid metabolic process (*APOE*, *SREBF1*), regulation of cellular senescence (*CDKN1A*, *IGF1R*), interferon (IFN)-γ signalling (*ICAM1*, *HLA-DPB1*), and transcription factors associated with IFN signalling (*IRF-4*, *IRF-1*, *NF-kB*) (**Fig. 1C, Table S2**). Instead, mRNAs encoded by gene sets associated with cell cycle (*NASP, DPF1*) and telomere organization (*USP7, XRN1*) were al depleted in PMS iNSCs (**Fig. 1D**). This is concordant with previous work identifying cellular senescence in the NSCs of individuals with PMS.^17,24^

**Figure 1.**
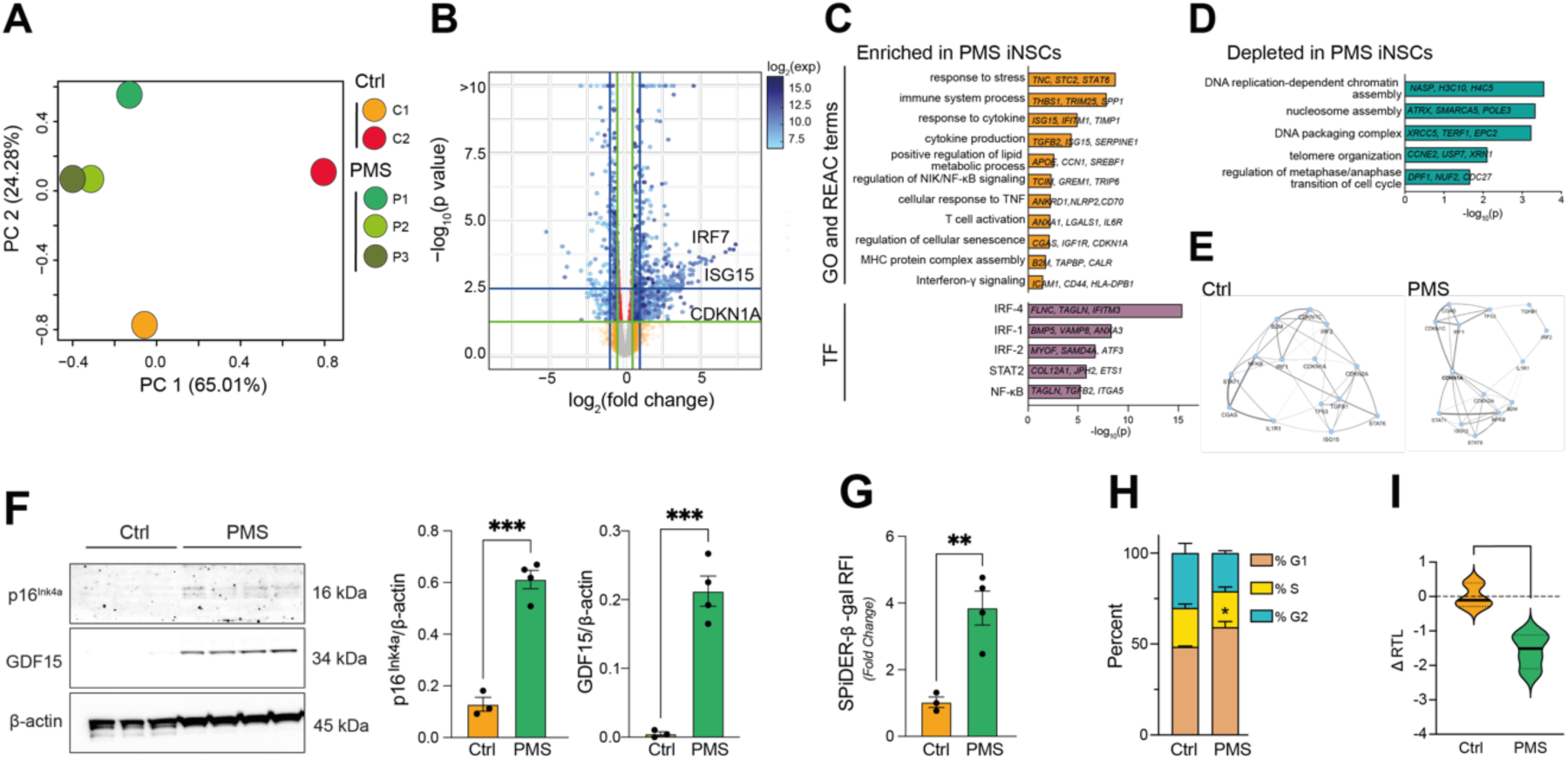
Bulk mRNAseq reveals increased inflammatory signalling and senescence markers in PMS iNSCs. **A)** PCA summarizing the co-variation of expression levels in the mRNA sequencing. C and P are independent cell lines (**Table S1**). (**B**) Volcano plot illustrating differentially expressed genes, *vs* log_2_ abundance in PMS iNSCs compared to Ctrl iNSCs. Log_2_(FC) *vs* adjusted p-values (with Benjamini Hochberg multiple testing correction) are reported. (**C-D**) Pathway enrichment analysis on GO and REAC terms and enriched transcription factors (TF) based on TransfFac annotation, from mRNA sequencing. (**E**) Gene regulatory networks, inferred on manually curated, differentially expressed genes contrasting of Ctrl *vs* PMS iNSCs networks, derived from the mRNA sequencing. (**F**) Representative western blots and quantification for p16^Ink4a,^ GDF15, and β-actin. (**G**) Quantification of relative fluorescence intensity (RFI) of senescence-associated β-galactosidase expression using SPiDER-β-gal. Data represented as a fold change over Ctrl iNSCs. (**H**) Flow cytometry-based quantification of iNSC cell cycle states. Data plotted as cells in percent of cell cycle state. (**I**) Quantification of changes in PMS relative telomere length (RTL) over Ctrl iNSCs. Experiments in **F-I** were done on n= 3 Ctrl and n= 4 PMS iNSC lines each performed in n= 3 replicates. Data in **F-I** are mean values ± SEM. *p ≤ 0.05, **p≤ 0.01, ***p≤ 0.001, un-paired t-test, with unequal variances.

Next, we investigated the dynamics of gene regulatory networks (GRNs), inferred on differentially expressed genes, associated with enriched terms (inflammation, regulation of cellular senescence, and interferon signalling). We observed a clear hub centred on the senescence gene *CDKN1A*, which then strongly interacted with *CDKN1C*, *CDKN2A*, *STAT1*, and *IRF1* in PMS iNSCs only (**Fig. 1E**). Previous studies have demonstrated that senescence induction via p21 (*CDKN1A*) and p16^Ink4a^ (*CDKN2A*) expression activates IFN response.^52^ Density plots of covariation of inflammatory and senescent transcripts (selected based on GO annotations) showed greater difference in weights (i.e. interaction strength/ co-variation in expression) between Ctrl and PMS samples (with stronger covariation in the PMS samples), compared with non-differentially expressed genes (**Fig. S1C**). These results support the predicted interactions between senescence and inflammation pathways.^52^ This GRN analysis also supports the hypothesis of an induction of a senescence program only in PMS iNSCs that then promotes IFN and inflammatory signalling activities.

Because the sequencing data showed upregulation of genes associated with cellular senescence, we independently validated these findings using Western blot and senescence-associated β-galactosidase (SA-β-gal) staining. We found upregulation of markers of cellular senescence p16^Ink4a^ and GDF15 (**Fig. 1F**)^53^ and increased expression of SA-β-gal using SPiDER-β-gal dye in all PMS lines (**Fig. 1G**).^54^ Cellular senescence is primarily associated with a halt in cell cycle, which was supported by a depletion of cell-cycle pathways in the mRNAseq (**Fig. 1D**). We then assessed the cycling stage of iNSCs using flow cytometry. Despite iNSCs being analysed under proliferative conditions in chemically-defined media, we still identified a significantly higher proportion of PMS iNSCs in the G1 phase of the cell cycle (*vs* Ctrl), which is typical of quiescent or senescent cells (**Fig. 1H**).^55^ Lastly, quantitative (q)PCR-based analysis of relative telomere length revealed that PMS iNSCs have significantly decreased telomere lengths (*vs* Ctrl) (**Fig. 1I**).

Therefore, PMS iNSCs phenotypically display intrinsic features of senescence, which are also reflected at the transcriptomic level via upregulation of pathways associated with inflammation and IFN signalling.

### PMS fibroblasts and iNSCs maintain pathological epigenetic hallmarks

We further investigated the potential origin of the senescent and inflammatory signatures and phenotypes seen in PMS iNSCs. We postulated that the direct conversion from fibroblasts into iNSCs would maintain or even facilitate the emergence of new epigenetic landscapes that further promote the senescence phenotype.

To this aim, we assessed the methylation status, genome-wide, using whole genome bisulfite sequencing (WGBS) on the parental fibroblasts and matched iNSC lines. PCA revealed a tight reproducibility of replicates and a clear separation between fibroblasts and iNSCs as well as between Ctrl and PMS samples (**Fig. 2A**).

**Figure 2.**
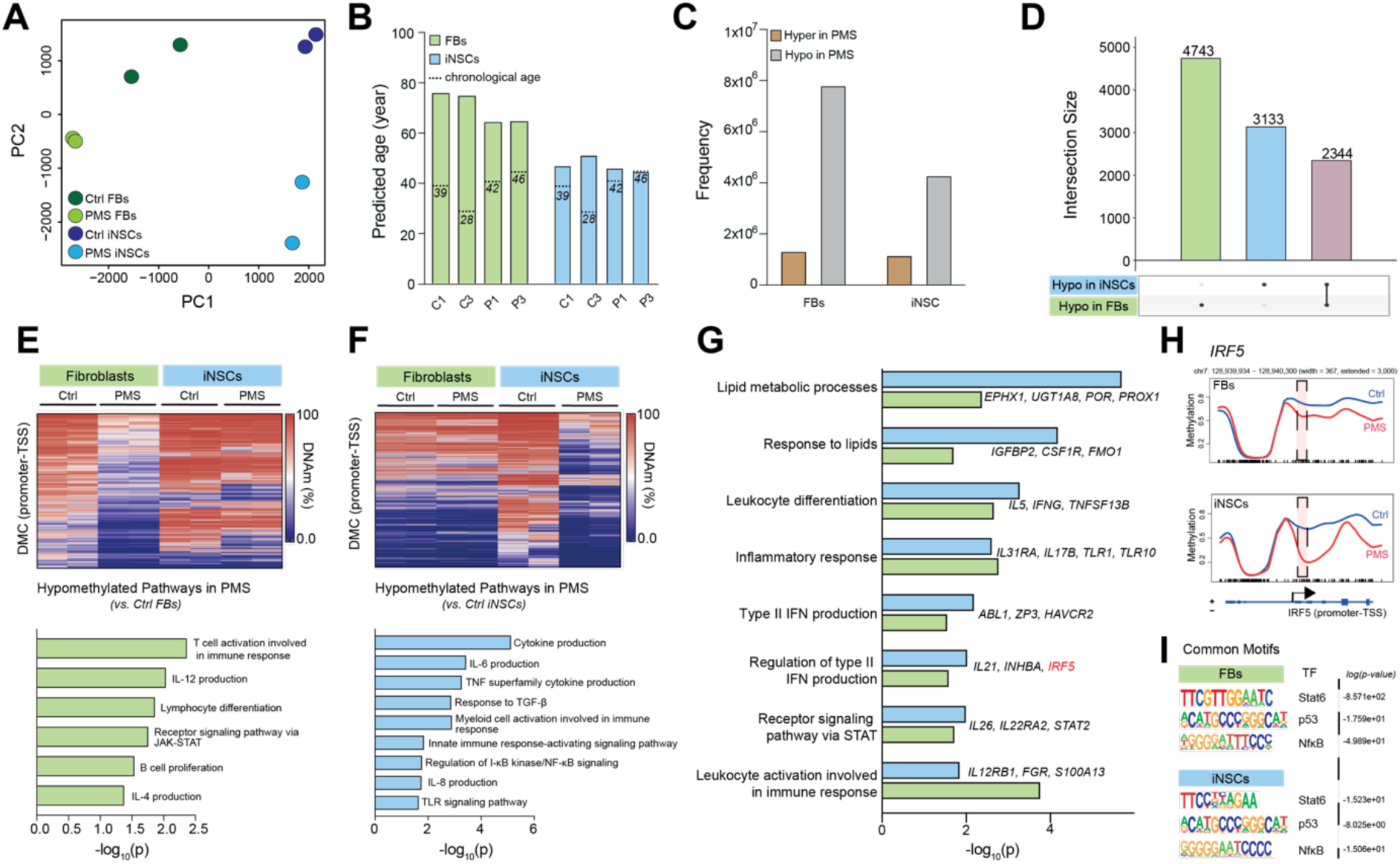
Whole genome bisulfite sequencing (WGBS) uncovers inflammatory pathways found commonly hypomethylated in PMS fibroblasts and iNSCs. (**A**) PCA summarizing methylation quantified in the WGBS data for fibroblasts and iNSC samples (**Table S1**). (**B**) Cortex age DNA methylation aging clock inference. Dashed line indicates chronological age at which FBs samples were sampled. (**C**) Frequency of differentially methylated cytosines (DMC), plotted as Ctrl *vs.* PMS. (**D**) UpSet plot of hypomethylated genes within the promoter-transcription start site (TSS) region. (**E-F**) Heatmap and enrichment analysis of hypomethylated genes. (**G**) Enrichment analysis of commonly hypomethylated genes in PMS (*vs* Ctrl) fibroblasts (light green) and iNSCs (light blue). (**H**) Example of methylation difference for *IRF5* (genome browser tracks), as in **G**. (**I**) Proportional sequence logos on HOMER motifs resulting from an enrichment analysis from **E**.

We identified 28 million CpG sites per sample and consistently observed increased hypermethylation in iNSCs (*vs* fibroblasts) (**Fig. S1D-E**). To provide insight to the extent the direct reprogramming reset the aging-related epigenome, we assessed the Cortex Age DNA methylation (DNAm) aging clock using methylation data from the human cortex.^56^ The cortex clock predicted an age similar to the chronological age of the donor cells used to generate the iNSCs, suggesting minimal epigenetic resetting occurred during the direct reprogramming (**Fig. 2B**). We also matched this modality against the Horvath and Zhang DNAm clocks, which further confirmed the maintenance of epigenetic age after direct reprogramming for most cell lines (**Fig. S1F**).^57,58^ Thus, our own direct reprogramming technology to generate iNSCs from skin-derived fibroblasts maintains epigenetic information from the donor cells.

We next investigated methylation commonalities and specific differences between Ctrl and PMS samples in both fibroblasts and iNSCs. We assessed the distribution of differentially methylated regions (DMRs) across genic and intergenic annotations, including a class linking the DMR to the transcription start sites (TSS) and the resulting distribution was quasi uniform (**Fig. S1G-H**). The differentially methylated cytosines (DMCs) and DMRs indicated an increased hypomethylation in both PMS fibroblasts and iNSCs (*vs* Ctrl; **Fig. 2C, Fig. S1I**). To further explore how these differences in methylation profiles account for transcriptional and phenotypic changes, we investigated genes with hypomethylated DMRs located in proximity of the TSSs. We identified 4,743 genes hypomethylated only in PMS fibroblasts, 3,133 genes hypomethylated only in PMS iNSCs, with 2,344 hypomethylated genes shared between PMS fibroblasts and iNSCs (**Fig. 2D**).

We performed GSEA on the specific hypomethylated genes of the PMS fibroblasts, and found pathways associated with T cell activation, IL-12 production, and JAK-STAT signalling (**Fig. 2E, Table S3**). This was further supported enriched motifs associated with transcription factors in the PMS fibroblasts. These include *SREBP1*, known to regulate T cell growth and survival, as well as *DDIT3*, which is closely connected to JAK-STAT signalling (**Fig. S1J**).^59,60^ Gene pathways specific to hypomethylation PMS iNSCs included cytokine production, TNF superfamily cytokine production, and regulation of I-κB kinase/NF-κB signalling (**Fig. 2F**). Additional analyses of enriched motifs identified *STAT5* and *IRF6*, encoding proteins known to be involved in cytokine production, immune response, and senescence (**Fig. S1J**),^61,62^ and *ARID5A*, which encodes a protein involved in the immune response by stabilizing *IL-6* mRNA (**Fig. 2F, Fig. S1J**).^63^

To determine which pathways were epigenetically modulated between the PMS fibroblasts and iNSCs, we performed GSEA on genes commonly hypomethylated in both cell types. The results revealed genes encoding proteins with functions in pathways associated with lipid metabolism, inflammation, and IFN production (**Fig. 2G**). In a separate study, we performed metabolomics and lipidomics on the same Ctrl and PMS cell lines, which led to the identification of increased cholesterol synthesis in PMS iNSCs and a new role for this pathway in establishing and sustaining their pathological and neurotoxic phenotype.^46^ In addition, using published GWAS studies, genes associated with MS progression and pathology were identified such as leukocyte activation and differentiation, STAT signalling, and IFN production.^64–66^ Another example is *IRF5*, known to have associated gene variants in MS,^67^ which we found hypomethylated at the promoter-TSS region, in both PMS fibroblasts and iNSCs (**Fig. 2H**).

We also analysed hypermethylated genes, with DMRs located in the promoter regions and in proximity of the TSS. We found 2,946 genes specifically hypermethylated in PMS fibroblasts (*vs* Ctrl), 858 specifically hypermethylated in PMS iNSCs (*vs* Ctrl), and 291 shared hypermethylated genes (PMS fibroblasts *vs* Ctr iNSCs) (**Fig. S1K**). Analysis of the unique differentially hypermethylated genes revealed differences in pathways associated with RNA metabolic processes and cell cycle in fibroblasts and pathways associated with transcription and neuronal differentiation in iNSCs (**Fig. S1L-M**). The hypermethylation pattern of genes that is shared between PMS fibroblasts and iNSCs included DNA-templated transcription and telomerase holoenzyme complex assembly (**Fig. S1N**).

To further investigate the epigenetic regulatory modules defined from the WGBS dataset, we used de novo and directed HOMER analysis is to assay for enrichment of shared binding motifs between the PMS fibroblasts and iNSCs. This analysis identified NF-κB, a major transcription factor that regulates genes responsible for both the innate and adaptive immune response and is associated with senescence (**Fig. 2I**).^66,68,69^ These results were corroborated by the GSEA of the mRNAseq data, where an enrichment in genes associated with NF-κB was also observed in the PMS iNSCs (**Fig. 1C**).

Our data suggest that PMS pathology is strongly linked to alterations of the epigenome, which we identified first in patient fibroblasts and confirmed in iNSCs. Many of these epigenetic differences are features of senescence and involve genes that regulate inflammatory, metabolic/lipid, and IFN pathways. Furthermore, when PMS fibroblasts are directly reprogrammed into iNSCs they also adopt an epigenetic landscape that is permissive for increased expression of proteins associated with secretion of inflammatory cytokines, specifically IL-6 and TNF-α.

### iNSCs share an RG-like signature that is identified in transcriptomic signatures from post-mortem human datasets

Next, we explored the heterogeneity of iNSCs and the respective subpopulations driving the phenotypes observed in mRNAseq and WGBSseq sequencing.

We first performed single cell (sc) and single-nucleus (sn) RNAseq coupled with ATAC sequencing on the Ctrl and PMS iNSCs to determine if there was a subpopulation of cells that were driving the phenotypes observed in the mRNAseq and WGBSseq (**Fig. S2A**). To minimize technical discrepancies between the two approaches, data-driven, specific filters were applied, on the proportions of reads incident to mitochondrial (MT) DNA and ribosomal proteins (RP). We retained cells with 15-40% RP ratios for the scRNAseq samples and 2-25% RP ratios for the snRNAseq data samples; across the dataset, a 20% MT ratio filter was applied (**Fig. S2B-C**). Additional filters rely on number of UMIs > 8,000, number of genes per cell > 1,000, log_10_ (genes per UMI) > 0.75. A total of 26,138 cells, across all samples, passed all filtering criteria, with median UMI counts per cell of 22,296. The average number of cells per sample was 3,267, ranging from 1,277 to 5,545; the average number of genes per cell was 5,725, ranging from 4,357 to 6,731 (**Fig. S2D-E**).

Using these filtering criteria for RNA analysis, we identified a total of 8 clusters (**Fig. 3A**). We applied the ClustAssess^70^ framework to determine the optimal parameters in a data-driven way using the Element Centric Similarity (ECS)^71^ as assessment criteria for the crisp partitioning of cells. The type of features (highly variable or most abundant) and number of features retaining signal (i.e. not biased by noise or shallow sequencing) were first determined on 20 iterations of ECS and summarized as element centric consistency (ECC). Next the number of neighbours for the community detection approach and the clustering method was also determined on high ECC distribution. The stable configurations linked number of clusters, number of the most frequent partition, and the resolution parameter (**Fig. S2F-G**). The distribution of ECC across the UMAP indicated a high stability for the selected number of clusters (**Fig. 3B**). We observed a quasi-uniform distribution of cells across all samples and conditions for all clusters (**Fig. S2H**).

**Figure 3.**
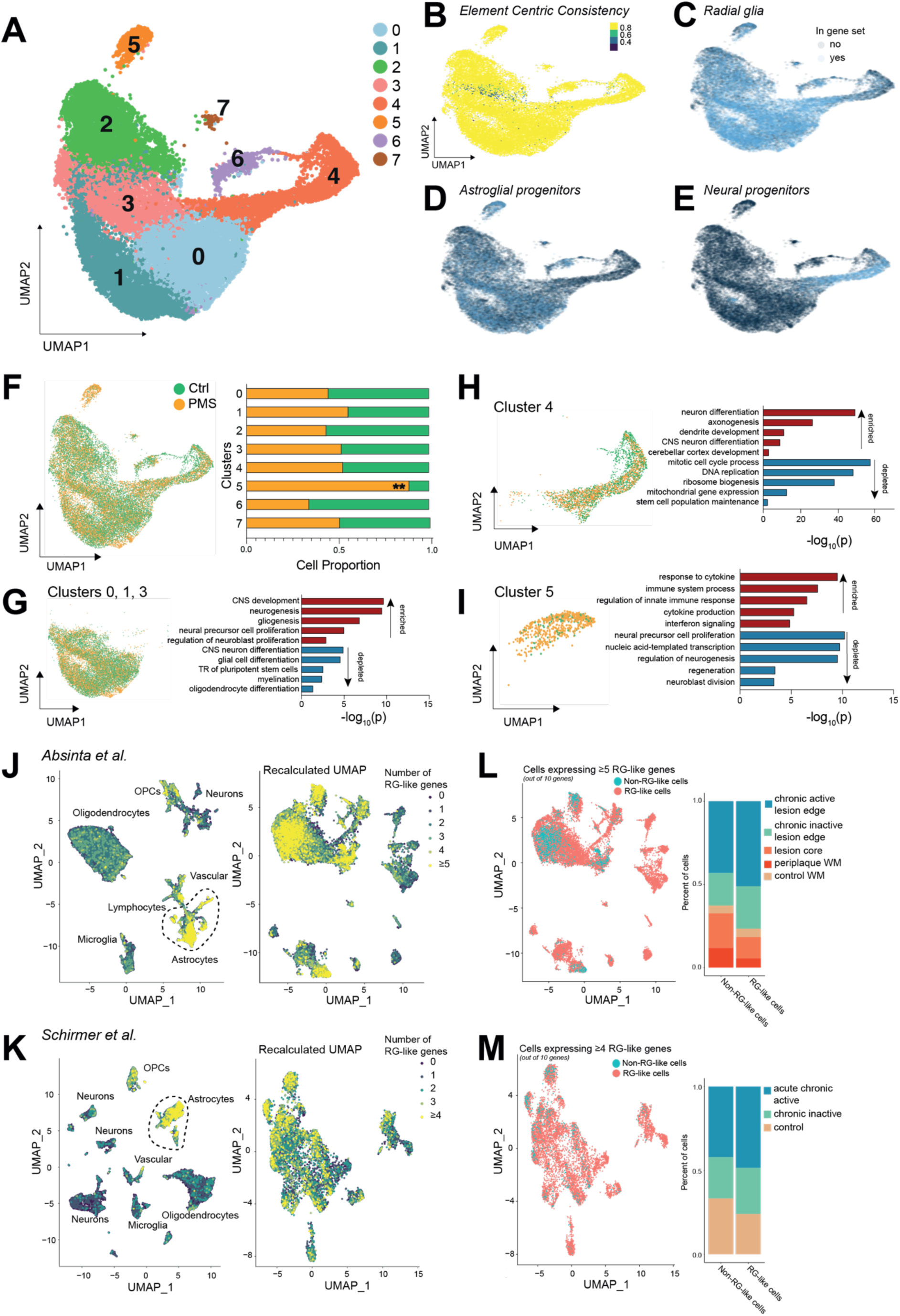
iNSCs display a RG-like transcriptomic signature that can be identified in the adult human brain using single nuclei sequencing. (**A**) UMAP of Ctrl and PMS iNSC single-cell, -nucleus RNAseq samples, post quality checking and filtering; a stable partitioning of cells is also highlighted. (**B**) Element centric consistency, derived on 30 iterations, calculated per cell, and visualized on the RNA UMAP. (**C-E**) Voting scheme of genes associated with a RG-like signature (**C**), astroglial progenitor signature (**D**), and neural progenitor signature (**E**) as in **A**. (**F**) Cluster distribution in human iNSCs on the RNA UMAP. The histogram summarizes cell proportions per cluster. **p ≤ 1e-106 (X^2^ test). (**G-I**) Enrichment analysis of enriched and depleted terms in clusters 0, 1, 3 (**G**); cluster 4 (**H**); cluster 5 (**I**) *vs* all other clusters; recalculated RNA UMAPs illustrating the distribution Ctrl *vs* PMS cells. (**J-K**) Recalculated UMAPs of RG-like cells in two *ex vivo* MS datasets, *Absinta et al., 2021* (**J**) and *Schirmer et al., 2019* (**K**). OPCs, oligodendrocyte progenitor cells. (**L-M**) Voting scheme UMAPs and histograms summarizing the frequency of RG-like cells (and non-RG-like cells per area of interest as in **J-K**. WM, white matter.

An indirect assignment of cluster identity was based on co-localisation of expression of standard genes for radial glia (RG), astroglial progenitors, and neuronal progenitors (gene lists in **Table S4**). We found that a majority of the iNSC clusters were defined by an RG-gene or astroglial progenitor gene signature, including the expression of *SOX2*, *NES*, *PAX6*, *PTPRZ1*, *HES1*, and *CKB* mRNAs (**Figs. 3C-D**). The majority of clusters (0-3, 5-7) had a radial glia and/or astroglial progenitor gene signature, encompassing 20-80% of cells within each individual cluster, defined expression thresholds (**Fig. S2I-J**). We also found a small proportion of neural progenitor cells primarily represented by cluster 4 (**Fig. 3E, Fig. S2K**). Genes associated with cell differentiation, including oligodendrocyte progenitor cells (*PDGRA*), oligodendrocytes (*OLIG1*, *OLIG2*, *MBP*), astrocytes (*AQP4*, *ALDH1L1*), and mature neurons (*CALB*, *CCK*) were lowly expressed across the samples and clusters (**Fig. S2L,** gene list in **Table S4**).^72–75^ This initial voting-scheme analysis suggests that proliferating iNSCs – similarly to hiPSC-NSCs^76^ – are a heterogenous population of cells displaying a transcriptional signature reminiscent of RG-like, astroglial progenitor cells, and a small subpopulation of neural progenitor cells, with little to no detection of terminally differentiated cells.

We next assessed the proportion of cells belonging to either Ctrl or PMS iNSCs within the individual clusters. Strikingly, we found mostly equal representation amongst all clusters but cluster 5, which was significantly enriched with PMS iNSCs (**Fig. 3F**). Towards further understanding the biological role of the individual clusters we performed GSEA. Core clusters 0, 1, and 3 were enriched for terms linked to CNS development, gliogenesis, and proliferation, while depleted of terms associated with cellular differentiation and pluripotency (**Fig. 3G, Table S5**). Cluster 4 was enriched in pathways related to neuronal and cortical development, with a coordinated depletion in the mitotic cell cycle genes (**Fig. 3H**). The remaining clusters were associated with mitochondrial organization and oxidative phosphorylation (cluster 2), glial cell differentiation (cluster 6), and ion transport (cluster 7) (**Fig. S3A-C**). Cluster 5, showing a striking 6-fold higher frequency in PMS iNSCs (85.6% PMS *vs* 14.4% Ctrl) (**Fig. 3F**), was characterized by genes enriched for cytokine production, immune processes, and IFN signalling, and genes depleted for proliferation and regeneration (**Fig. 3I**).

To evaluate the contribution of cell cycle genes to the transcriptomics signature of clusters, we identified the cell cycle stage using *a priori* defined gene sets. We note a strong representation of cells expressing G1-phase specific genes in cluster 4, a depletion of cells expressing G1-phase specific genes in clusters 1, 2, 3, and a depletion of cells expressing S-phase specific genes in cluster 5, which further supports the GSEA analysis per cluster (**Fig. S3D**).

To investigate the relevance of the *in vitro* iNSC model to human disease, we aligned our *in vitro* results with two independent, publicly available *ex vivo* human snRNAseq datasets from post mortem MS cases and controls.^16,77^ Using a panel of canonical RG genes that were also used to characterize the iNSCs (**Table S4**), we identified disease associated RG-like cells within the annotated astrocyte clusters in both datasets (ranging between 6.5 – 7.8% of total astrocyte cluster, **Fig. 3J-K**). When compared to the non-RG-like cells within the astrocyte cluster, RG-like cells exhibited a significantly higher proportion and expression of RG genes *ETNPPL*, *PTPRZ1*, *SOX2*, *PAX6*, and *PCNA* encoding a cell cycle marker (**Fig. S3E-F**). RG-like cells expressed astroglial genes (**Fig. 3C-D**) and very little microglia-specific or oligodendrocyte progenitor cell-specific genes (**Fig. S3E-F**), further supporting their identity. To determine whether these newly identified RG-like cells hold neurogenic potential, we assessed related genes, including *SOX11*, *DCX*, and *TUBB3* and found little expression in both RG-like and non-RG-like cells within the astrocyte cluster in both datasets (**Fig. S3G**). A large proportion of the RG-like cells expressed genes specific of G2M or S phases, which indicated their ability to progress through the cell cycle (**Fig. S3H**). Lastly, we assessed the proportion of RG-like cells across the different MS lesion types. Out of all the RG-like cells in both datasets, we identified that 50% were in chronic active lesions, whereas the smallest proportion of cells were found in control tissue (**Fig. 3L-M**).

Therefore, we identify a small proportion of non-neurogenic RG-like cells in the healthy adult human brain, which significantly increase in frequency in chronic active lesions in the PMS brain.

### Patient iNSCs harbour a senescent, IFN-responsive RG-like cell cluster reminiscent of Disease Associated RG the PMS brain

As cluster 5 was predominant in PMS iNSCs (*vs* Ctrl), we sought to further the disease-associated transcriptomic signature of this cluster. Transcriptionally across all samples, cluster 5 showed a significant enrichment of genes associated with cellular senescence, IFN α/β signalling, and RIG-I signalling, along with a depletion of genes associated with cell proliferation, DNA-templated transcription, and NOTCH1 signalling (**Fig. 4A**). Cluster 5 also had the highest expression of genes associated with IFN-α and -γ response and the SenMayo gene set^78^ (*vs* core clusters 0-3) (**Fig. 4B**). We performed a differential expression analysis followed by GSEA of cluster 5 only specific markers and identified a strong enrichment for IFN and cytokine signalling pathways and SASP that was associated with high expression of *IFIT1*, *ISG15*, and *NLRP2* in PMS iNSCs (*vs* Ctrl) (**Fig. 4C**). We confirmed the high expression of IFN-response genes (*IFIT1*, *IFIT2*) was linked to the hypomethylated promoter regions associated with IFN signalling seen in both PMS fibroblasts and iNSCs, identified in WGBS analysis (**Fig. 2H**).

**Figure 4.**
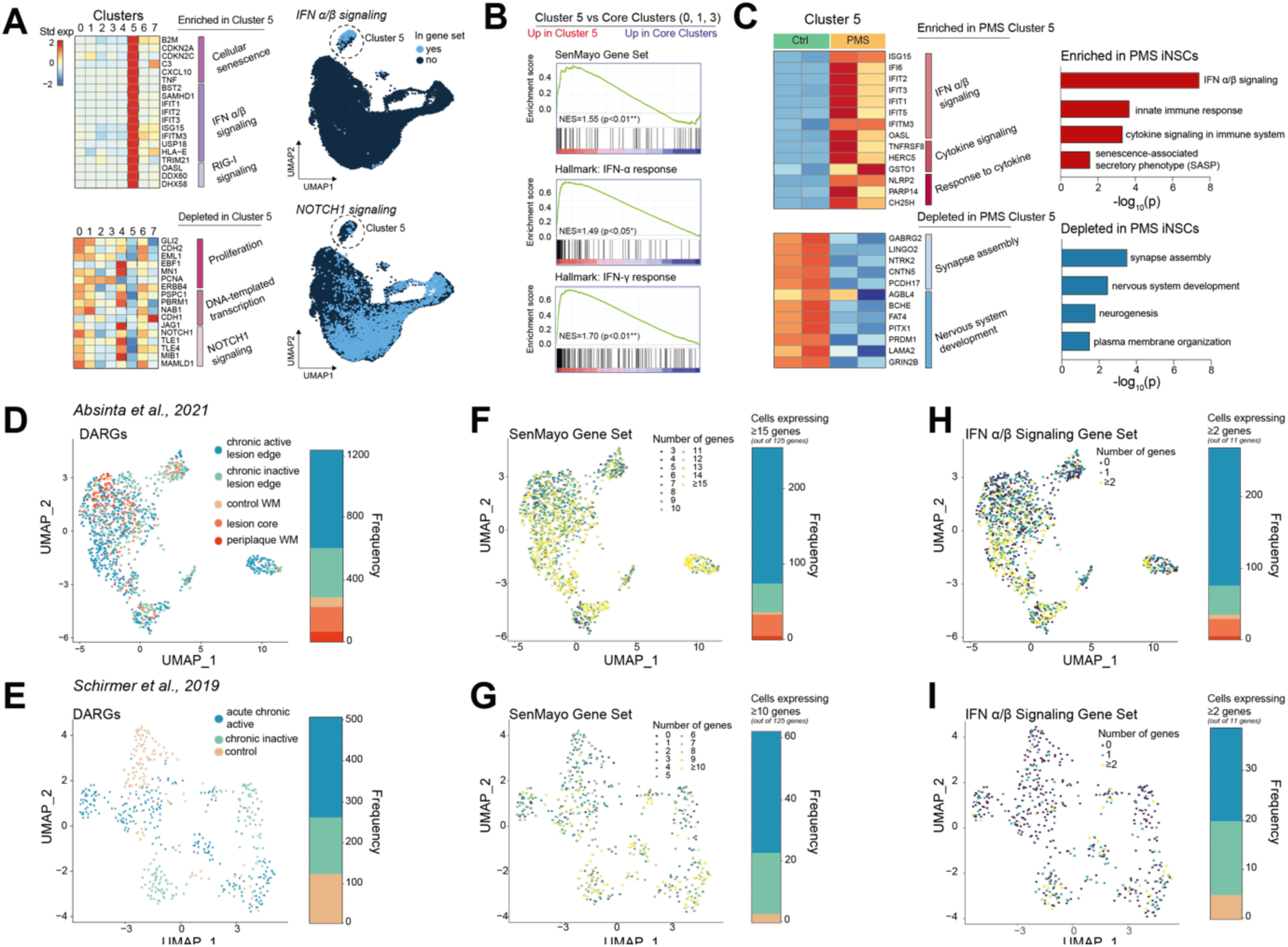
A specific PMS iNSC cluster displays senescence and IFN-signalling which is also identified in DARGs of the PMS brain. (**A**) Heatmap of RNA cluster 5 signature markers and summary of associated enrichment terms. Voting scheme UMAP of genes associated with IFN-α/μ signalling and NOTCH1 signalling. (**B**) Enrichment plots of the SenMayo gene set, and the IFN-α, and IFN-γ response in cluster 5. (**C**) Heatmap of standardized expression of highly expressed *vs* depleted transcripts, specific to only PMS cells *vs* Ctrl cells in RNA cluster 5 and selected enriched pathways significant for the selected genes. (**D-E**) Recalculated UMAP on the filtered cells identified as DARGs in Figure 3L-M. Stacked Histograms summarizing the frequency of cells per lesion area. (**F-I**) UMAPs and histograms of DARG frequency that express genes in the SenMayo gene set and IFN-α/μ signalling gene set respectively from *Absinta et al., 2021* and *Schirmer et al., 2019* data.

We next wanted to assess the expression of SenMayo and IFN α/β signalling gene sets in RG-like cells from the two human *ex vivo* snRNAseq datasets^16,77^ of post-mortem MS brains (**Fig. 4D-E**). A proportion (16-28%) of total RG-like cells – which we termed Disease Associated RG (DARGs) – in chronic lesions showed non-zero expression of the SenMayo gene set, whereas <5% of RG-like cells with the same features were identified in control tissues (**Fig. 4F-G**, **Fig. S4A-B**). We also identified an enrichment in IFN-associated mRNAs in DARGs located in chronic active lesions (**Fig. 4H-I, Fig. S4C-D**).

We assessed whether the senescence and IFN-associated expression signatures are unique to DARGs by applying the same expression thresholds to the non-RG-like cells and analysing the datasets. Within chronic active lesions we identified a ∼2.3-fold increase in the proportion of senescent DARGs (*vs* senescent non-RG-like cells) in both datasets^16,77^ (**Fig. S4A-B**). We then compared all lesion areas and identified a ∼2-fold increase in the fraction of senescent DARGs in the edge of chronic active lesions, when compared to lesion core, chronic inactive lesion edge, periplaque white matter, and control white matter^16,77^ (**Fig. S4A-B**). DARGs in *Absinta et al.*^16^ showed high expression of IFN-associated genes (*vs* non-RG-like cells) in chronic active lesions (**Fig. S4C**), while displaying the same trend in chronic inactive lesions in *Schirmer et al.*^77^ (**Fig. S4D**).

These findings provide further support to the existence of non-neurogenic DARGs in the PMS brain, particularly in chronic active lesions, with an inflammatory and senescent transcription signature. Notably, the direct reprogramming of patient somatic cells into stably expandable iNSCs allows for the recapitulation of distinctive disease-associated cellular phenotypes and gene signatures found in the post-mortem MS brain.

### Disease associated senescent RG-like cells spread dysfunctional features towards other clusters

We then performed pseudotime analysis to better understand the developmental trajectories of the RG-like cell cluster of PMS iNSCs with senescent and inflammatory signatures (**Fig. 5**). We removed the neural progenitor-associated cells (cluster 4) which allowed our analysis to focus on the establishment of inflammatory cluster 5. The initialization of the pseudotime focused on cluster 5, defined as the endpoint. The predicted trajectory started with core clusters 0, 1, and 3, progressed to cluster 2 and ended in cluster 5 (**Fig. 5A**). A community-based clustering was applied on the gene expression levels, with the stability assessed using ClustAssess.^70^ Several gene modules i.e. clusters of genes with similar expression profiles across the pseudotime were predicted (**Fig. S4E**). We focused on three gene modules with distinct expression patterns. The expression profile of the first module (module 10) focused on the core clusters 0, 1, and 3. A gene enrichment analysis identified significantly elevated expression of genes associated with cell cycle terms, as well as TFs known to maintain NSC/RG identity such as *SP2* (**Fig. 5B, Table S6**).^79^ Module 3, which consisted of mostly cells assigned to cluster 2, showed enrichment in terms associated with mitochondria and antigen processing and presentation, including NSC/RG-associated TFs *E2F1* and *PAX6* (**Fig. 5C**). These results support the GSEA enrichment on cluster 2 characterized by mitochondrial and metabolic gene pathways (**Fig. S3B**). Module 7 mostly overlapped with cells in cluster 5 and exhibited enrichment in IFN and cytokine signalling pathways, as well as TFs associated with IFN signalling (*IRF3*, *STAT2*) (**Fig. 5D**). This suggests that the progression towards cluster 5 may originate in iNSCs with a cluster 2-like gene signature and display pathways associated with mitochondria and cellular metabolism. Therefore, an altered metabolic signature in PMS iNSCs, which we have recently described,^46^ may promote the resurgence of the newly identified IFN responsive RG-like cell cluster 5.

**Figure 5.**
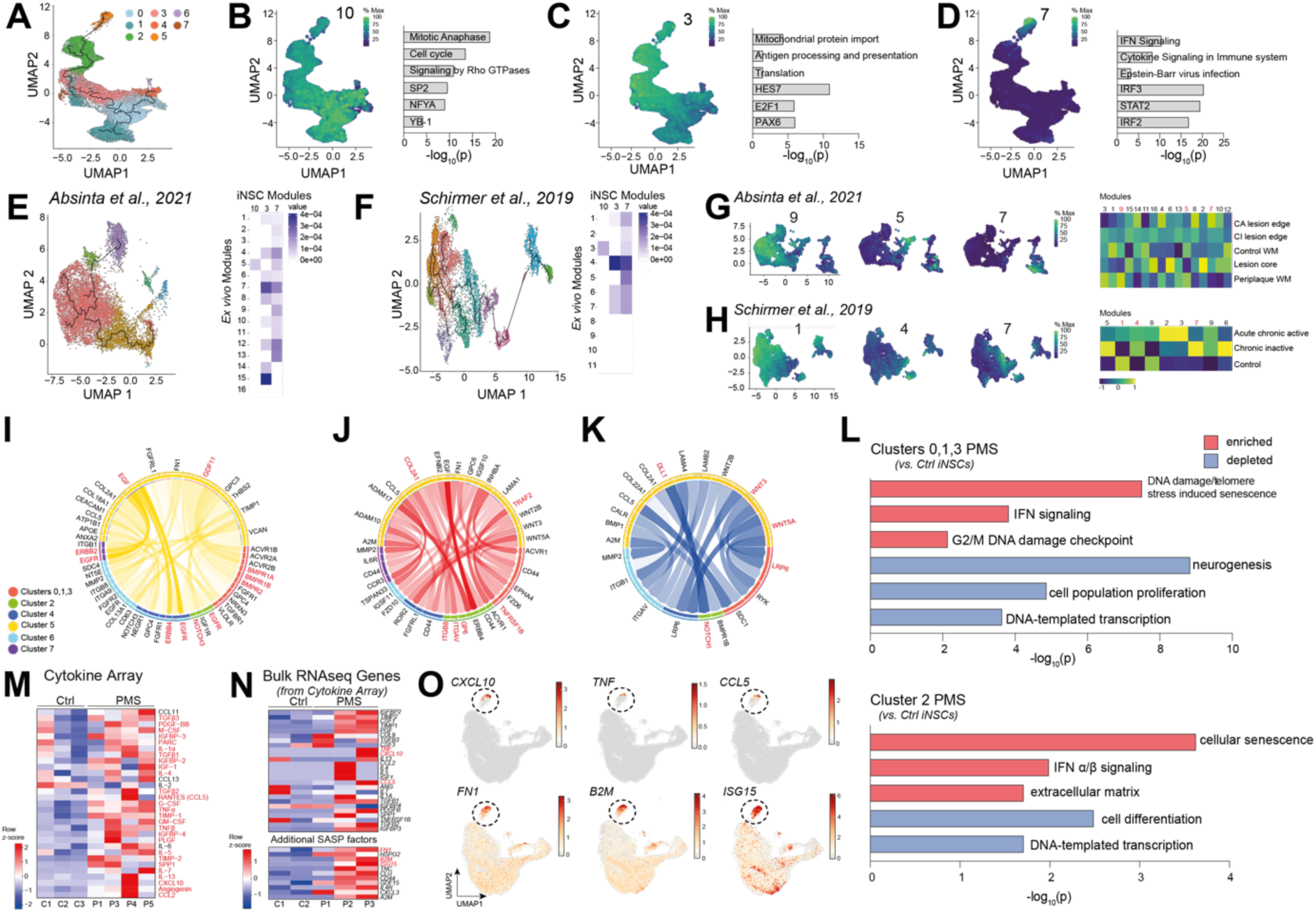
PMS iNSCs secrete a pro-inflammatory SASP that induces upregulation of genes associated with inflammation and senescence. (**A**) Pseudotime trajectory inferred and displayed on the *in vitro* dataset UMAP. Cluster 5 was used as initialization point (*endpoint*). (**B-D**) UMAP of the distribution of intensity of genes clustered in modules 10 (**B**), 3 (**C**), and 7 (**D**), determined based on the pseudotime ordering. GSEA summary and TFs corresponding to gene modules 10 (**B**), 3 (**C**), and 7 (**D**). (**E-F**) UMAP of pseudotime trajectory inferred from the *Absinta et al., 2021* (**E**) and *Schirmer et al., 2021* (**F**) datasets, respectively. The heatmaps summarize the scaled proportions of common genes matching between iNSC *in vitro* modules 10, 3, and 7 with *ex vivo* modules. (**G-H**) Modules selected from the *ex vivo* datasets as in **E-F** recapitulating the manually curated modules (**B-D**) identified on the *in vitro* pseudotime trajectory. The heatmaps summarize the scaled averaged expression of the *in vitro* gene modules, projected on the *ex vivo* datasets. (**I-K**) Circos plots of the intercellular ligand-receptor interactions predicted using NicheNet. (**I**) Yellow directed edges indicate interactions, (**J**) red edges summarize enriched ligand/receptor interactions i.e. upregulated target genes, (**K**) blue edges summarize depleted ligand/receptor interactions i.e. downregulated target genes, between cluster 5 and complement clusters in Ctrl and PMS iNSCs. (**L**) Enrichment summary on enriched and depleted genes in clusters 0,1,3 and cluster 2 (Ctrl *vs* PMS). (**M**) Heatmap of cytokine array performed on CM. Colour intensities are proportional with standardized normalized intensities. Proteins investigated also in the mRNAseq (**N**) are highlighted in red. (**N**) Heatmap of standardized normalized expression levels of genes coding for secreted proteins as in **M**. Genes investigated also in the single cell data (**O**) are highlighted in red. (**O**) Expression gradient UMAPs of the selected secreted proteins as genes (**N**).

Using the modules derived from the *in vitro* dataset, we next identified cells with similar transcriptomic signatures in the *ex vivo* post-mortem datasets.^16,77^ Focusing on the recalculated *ex vivo* UMAPs and underlining the RG-like cells (**Fig. 3J-K**), we inferred pseudotime trajectories on both datasets using the core of the RG-like cells (clusters 0, 1, 3) as an initialization point. (**Fig. 5E-F**). Next, we matched the gene modules determined in the *in vitro* and *ex vivo* datasets, respectively. Using the gene lists from the individual modules we cross-referenced the gene modules from the *in vitro* perspective (**Fig. 5B-D**, **Table S6**) and from the *ex vivo* perspective (**Fig. S4F-G**). For the *Absinta et al.* dataset we found coordinated gene expression within modules 5 and 7 that matched our *in vitro* curated modules and module 7 matched the inflammatory, senescent cluster 5 (**Fig. 5E, G**) and was associated with DARGs in the chronic active lesion (**Fig. 5G**). In the *Schirmer et al.* dataset we found coordinated gene expression within modules 4 and 7 (**Fig. 5F, H**). Genes that strongly contributed to module 7 were associated with chronic inactive lesions (**Fig. 5H**).

This analysis highlights the identification of a new cluster of senescent-like, inflammatory non-neurogenic DARGs stemming from astrocyte-like cells in the post-mortem MS brain.

As senescence is associated with and perpetuated by secreted proteins, we next investigated ligand-receptor interactions between cluster 5 (as source of ligands) and other clusters (as source of receptors) in iNSCs using NicheNet^80^. With this modelling, Ctrl iNSCs, were enriched for ligand-receptor interactions regulating cell maintenance and differentiation (i.e., Notch signalling) (**Fig. 5I**).^81^ However in PMS iNSC, modelling predicted strong interactions between TRAF2 in cluster 5 with TNFRSF1B in clusters 0, 1, and 3, which anticipates induction of NFκB activation^82^ and NSC activation,^83^ along with a depletion in Wnt signalling via LRP6 (**Fig. 5J-K**).

Next, we performed a cluster-by-cluster GSEA analysis and found that clusters 0, 1, and 3 in PMS iNSCS were enriched for senescence pathways, particularly those associated with DNA damage and corresponding depleted for proliferation-related terms (**Fig. 5L, Table S7**). We also detected significant interactions between *COL2A1* from cluster 5 and integrin-based receptors in cluster 2 (*ITGB8*, *ITGAV*, *GP6*), coupled with a depletion in NOTCH1 signalling (**Fig. 5J-K**). Enrichment analysis of genes in cluster 2 of PMS iNSCs further indicated enrichment in senescence, IFN signalling, and ECM and corresponding depletion in differentiation and DNA transcription that is known to be regulated by NOTCH signalling (**Fig. 5L**). Analysis of clusters 4, 6, and 7 consistently identified enrichment of inflammatory-associated terms (i.e., neurodegeneration, oxidative stress-induced senescence, and signalling by Ils) and a depletion in terms associated with NSC maintenance (i.e., Wnt signalling and cell cycle) (**Fig. S4H**). The enrichment in inflammatory terms in PMS iNSCs were linked to increased inflammatory interactions, stemming from cluster 5, between CCR3-CCL5 and between A2M-MMP2 (**Fig. 5J**).

To further test the hypothesis that the ligand-receptor interactions predicted above are relevant, we ran a cytokine array on the conditioned media (CM) from the bulk iNSC lines. Quantification of the cytokine array confirmed an increased secretion of cytokines associated with the SASP^84^ and inflammation (IL-6, IGFBP-3, and TNFα) in PMS iNSCs (*vs* Ctrl) (**Fig. 5M**).

We then quantified the expression levels of the genes coding for the upregulated SASP (**Fig. 5M**) using our mRNAseq data (**Fig. 1A-E**). Both *TIMP2* and *IGFBP2*, along with the known SASP gene *GDF15*, were upregulated both at gene and protein levels (**Fig. 1F** and **Fig. 5M-N**). Next, we re-analysed the single-cell data for the genes and proteins associated with the secreted factors identified in the mRNAseq and cytokine array, respectively, and confirmed the elevated expression in cluster 5 of *TNF*, *FN1*, and *ISG15* (**Fig. 5O**).

Overall, our findings suggest that the developmental trajectories of RG-like cells in cluster 5 arise from cluster 2. Additionally, PMS iNSCs secrete inflammatory factors as part of their SASP, and that this may induce a dysfunctional, senescent phenotype in cells in other clusters.

### Integrative multi-omics reveals regulons defining inflammatory RG-like cell in PMS iNSCs

To further investigate the epigenetic mechanisms that may contribute to the transcriptomic signature of the PMS iNSC cluster 5, we integrated the RNAseq data with single-nuclei chromatin accessibility data (snATACseq) using both matched and un-matched samples. For a high proportion of cells, the matched RNA and ATAC quantification was distributed proportionally across clusters (**Fig. S5A**). Next, we selected data-driven parameters and stable configuration (high ECC scores) on the ATAC modality using 30 iterations (**Fig S5B-D)** of ClustAssess. We identified 8 stable clusters on the ATACseq dataset.

We then projected the RNA clusters onto the ATAC UMAP, and the ATAC clusters onto the RNA UMAP, respectively to evaluate the concordance of the two modalities (**Fig. 6A-B**). Next, we quantified the equivalence between the RNA and ATAC clusters by summarizing matching cell types/states, which allowed highlighting the agreement between RNA cluster 5 and ATAC cluster 8, well separated in the non-linear space from the other cells (**Fig. 6C**). Then, we confirmed that the inflammatory ATAC cluster 8 was primarily composed of PMS iNSCs (**Fig. S5E**).

**Figure 6.**
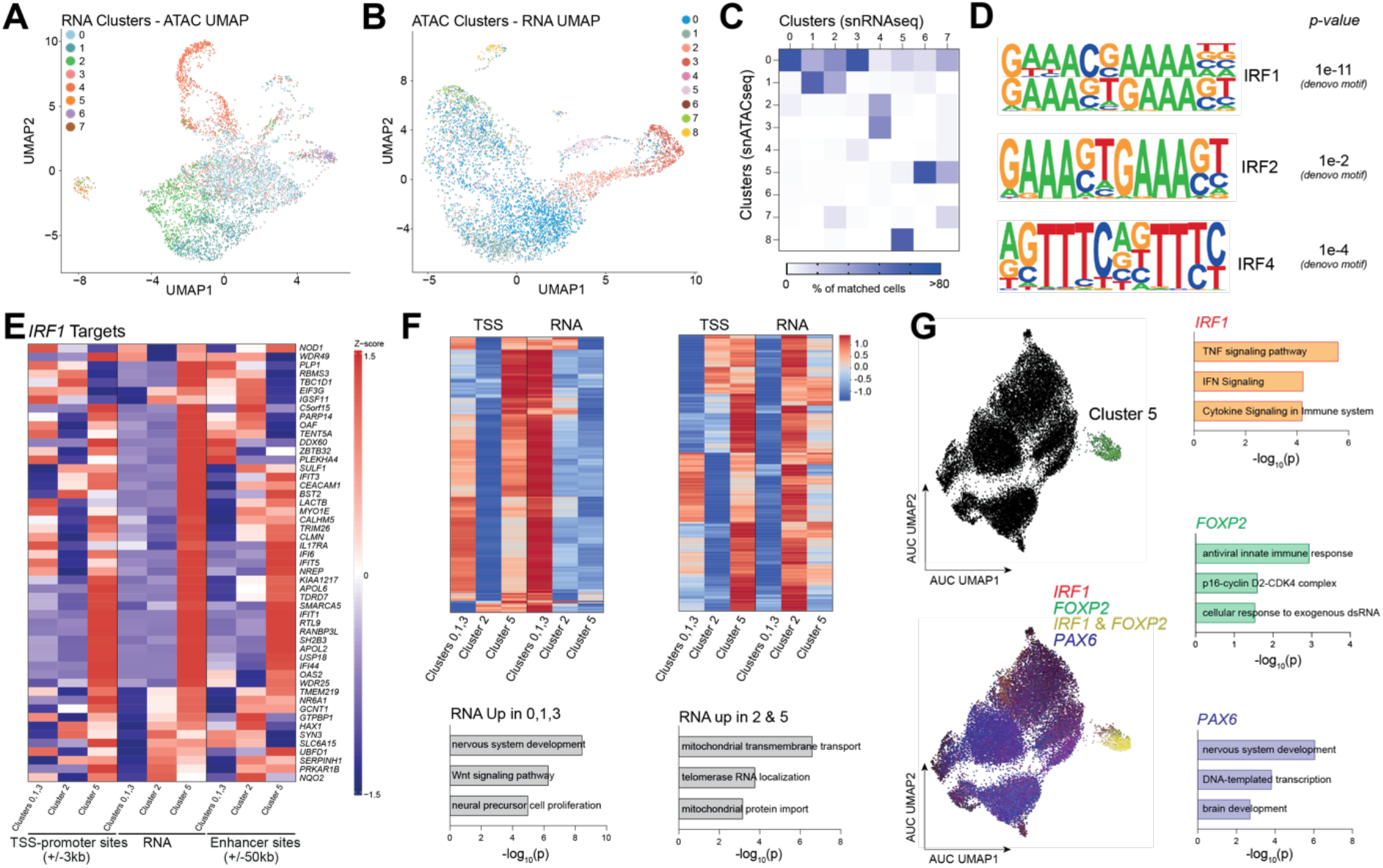
Multi-omics RNA/ATAC integration with further epigenetic characterization of cluster 5 cells. (**A**) ATAC UMAP of the overlaid localization of RNAseq clusters. (**B**) RNA UMAP of the overlaid localization of snATACseq clusters. (**C**) Heatmap of the percentage of matched assignations of cells across the snRNAseq and snATACseq clusters. The analysis was performed solely on matched cells i.e. cells with both RNA and ATAC expression. (**D**) Enriched motifs of signature and differentially accessible genes identified on the RNA cluster 5, and ATAC cluster 8. (**E**) Heatmap of accessibility (ATAC) and expression (RNA) of *IRF1* targets, i.e. TSS, RNA, and distal portions of marker genes identified as differentially expressed between grouped RNA clusters 0, 1, 3 *vs* cluster 2, and cluster 5, respectively. (**F**) Heatmap of marker genes, between clusters 0, 1, 3 *vs* 2 and 5. The summary of enrichment analysis applied on the selected genes. (**G**) SCENIC GRN inference summary and selection of IRF1 specific regulons, corroborated with enrichment analysis of regulons (*IRF1*, *FOXP2*, *PAX6*). The GRN inference was performed on the scRNAseq dataset.

We also examined differentially accessible regions (DARs) specific to inflammatory ATAC cluster 8, by selecting the corresponding peaks and enriched motifs on downstream genes, associated with TFs. Our analysis identified that DARs that gained accessibility were associated with immune processes, IFN signalling, and cytokine production (*IRF3*, *STAT2*), and DARs that lost accessibility were associated with genes pertaining to neuron and astrocyte differentiation and neural crest cell fate specification (*SOX4*, *SOX8*) (**Fig. S5F, Table S8**).

We next assessed the overlapping motifs significantly enriched for epigenetic changes in both PMS fibroblasts and iNSCs from the WGBS and from the predicted signature genes of RNA cluster 5, and identified common TFs including *p53*, *E2A*, and *SMAD2* (**Fig. S5G**). As *p53* and *E2A* are implicated in promoting immune function, involved in senescence, and in mediating IFN responses,^85,86^ our findings suggest that the chromatin accessibility for RNA cluster 5 closely predicts its RNA expression signature. Biologically, these cells are strongly IFN-responsive and display RG-like signatures. Our identification of common motifs maintained from the fibroblasts during reprogramming to iNSCs in PMS cell types further confirms the involvement of epigenetic regulation in perpetuating the senescent and IFN-response in the iNSCs.

Since IFN signalling was found to be strongly involved in RNA cluster 5, we next investigated the chromatin accessibility at the promotor regions. We identified increased accessibility of *IFIT1* in ATAC cluster 8, corresponding to increased expression in RNA cluster 5 (**Fig. S5H**). We next investigated genes that are known targets of the regulator of *IFIT1*, *IRF1*, which we identified to be enriched in the bulk mRNAseq of PMS iNSCs (**Fig. 1C**). *IRF1* is a key transcription factor implicated in facilitating TNF-α-induced senescence and is known to be anti-proliferative and pro-inflammatory.^87^ Within the IRF1 targets, notably *IFIT3*, *IFIT5*, and *OAS2*, known as IFN-response genes, we found clear associations between RNA expression (RNA cluster 5) and chromatin accessibility near TSS (+/− 3 kb) as well as in potential regulatory regions (+/− 50 kb from TSS) in ATAC cluster 8 (**Fig. 6E**). These data support the hypothesis that cells in RNA cluster 5 have permissive chromatin that underlies persistent activation of IFN-responses via IRF1.

The pseudotime analysis predicts RNA cluster 2 was most closely related, and strongly interacting with RNA cluster 5 based on ligand-receptor inference, so we next examined if a similar gene expression and chromatin accessibility signature could be identified in both clusters.

Some overlapping genes were found in cluster 2 with similar changes in nearby chromatin accessibility and included inflammatory genes *HAX1* and *SERPINH1* (**Fig. 6E**). Grouped clusters 0, 1, 3 exhibited little expression of *IRF1* target genes. GSEA on shared ATAC signatures, both for gain and loss of accessibility suggested that common gain of accessible regions were associated with stress response and blood brain barrier (BBB) maintenance, whereas loss of accessible sites correlates with notch and BMP signalling, and senescence pertaining to cell proliferation (**Fig. S5I-J**). This downregulation in notch and BMP signalling in cluster 2, based on interactions with cluster 5, was also predicted from the ligand-receptor interaction analysis (**Fig. 5K**). Importantly, not all changes in chromatin accessibility correlate directly to variation in mRNA (**Fig. S5J**).

To further understand patterns based on chromatin accessibility and RNA expression, we subdivided gene sets based on RNA upregulation in RNA clusters 0, 1, 3, and depletion in RNA clusters 2 and 5, and found enrichment for terms pertaining to Wnt signalling and NSC maintenance (**Fig. 6F**). We also found increased accessibility near the TSS of the RNA cluster 5 upregulated genes, again suggesting aberrant permissive chromatin in these cells promoting increased expression. When assessing coordinated upregulation of gene expression in RNA clusters 2 and 5, we found enrichment in terms associated with mitochondrial transmembrane transport, protein import, and telomerase RNA localization (**Fig. 6F**). These results support a close interaction between RNA clusters 2 and 5, with cluster 5 being specifically IFN-responsive.

Lastly, we inferred single-cell regulatory networks using SCENIC^88^. We observed that RNA cluster 5 was defined by two major regulons, *IRF1* and *FOXP2*, that was associated with genes regulating TNF and IFN signalling, as well as the p16-cyclin complex related to senescence (**Fig. 6F, Table S9**). *PAX6* was also identified as a major regulon across all clusters of the iNSCs and was defined by genes associated with nervous system development, supporting their RG-like state. Overall, RNA cluster 5, primarily represented in PMS iNSCs, is consistently defined by gene regulatory patterns that are associated with an IFN-responsive and senescence state.

In summary, our work demonstrates that direct reprogramming technology maintains hallmarks of PMS in cells due to maintenance of epigenetic memory. We reveal that patient fibroblasts have hypomethylation at genes associated with lipid metabolic processes and IFN signalling, which then became further accentuated upon direct induction into iNSCs.

Within the heterogeneous PMS iNSCs, we identify a novel disease associated cluster of IFN-responsive, inflammatory RG-like cells that display senescent features and are regulated by IRF1, which may spread dysfunctional features towards other clusters through their secreted factors.

Integration with publicly available datasets further identifies and highlights a long-neglected non-neurogenic DARG population, which is found significantly increased in chronic active lesions areas and display IFN and senescence gene expression.

## DISCUSSION

PMS is a complex neuroinflammatory and neurodegenerative disease that results from the interaction between environmental factors and genetic predisposition. The majority of genetic risk factors that have been identified in the development and progression of MS are associated with the peripheral immune system, mapping mainly to T cells, however recent work has also uncovered variants in genes expressed by glial cells.^89,90^ MS severity has also been linked to variants involved in mitochondrial function, synaptic plasticity, and cellular senescence in genes expressed in the CNS.^18^ To further understand how intrinsic glial cell dysfunction in PMS contributes to disease pathology we generated directly reprogrammed iNSCs.

Characterization of directly reprogrammed iNSCs showed PMS donor-derived cells reproduced reported phenotypes of increased expression of inflammatory signalling and senescence associated genes compared to Ctrl cells.^17^ Further, patient iNSCs maintained shorter telomere lengths compared to controls, indicating this type of reprogramming better maintains features of the donor cells that may be critical in driving the PMS phenotypes. We then established that iNSCs also maintained epigenetic signatures, as determined by DNA methylation age, after reprogramming from fibroblasts, indicating that this method of cellular reprogramming may conserve epigenetic information important for disease phenotypes. Globally, cell lines derived from people with PMS were found to have lower levels of DNA methylation, with enrichment of hypomethylation at promoter regions of targets for inflammatory transcription factors, such as STAT6 and NFκB. Interestingly, global hypomethylation has been identified to occur with aging in various organs in both mice and humans, which is believed to contribute to genome instability.^91^

Concordantly, hypomethylated regions in the PMS fibroblasts and iNSCs were enriched for pathways involved in immune response, suggesting global aberrant epigenetic regulation in individuals with the disease. Previous work has characterized global methylation signatures in the whole blood of people with MS and identified that half of the differentially methylated positions mapped to genes enriched in CNS cells and pathways.^92^ This work identified neurodegenerative-related pathways as epigenetically dysregulated in severe MS cases, which correlated with acceleration of methylation age.^92^ Intriguingly, we also characterize a loss of methylation in genes that regulate lipid metabolism, implicated in Ionescu, Nicaise et al.^46^ Given these pathways appear to be highly relevant to PMS phenotypes, we believe human iNSC methodologies provides an excellent, and biologically valuable, platform for studying mechanisms driving disease.

Combined single-cell and single-nucleus RNA data analysis allowed us to better characterize heterogenous iNSCs. Here we identified cells with both neural and RG phenotypes, and importantly were able to find cells with matching transcriptional profiles from datasets generated from post-mortem MS brains. We found that most of the cell clusters were identified by radial glial genes (*SOX2*, *PAX6*), with one cluster which we identified with a neural progenitor signature (*DCX*, *ASCL1*). Within these clusters, we define a novel subset of cells that is predominantly composed of cells derived from PMS donor fibroblasts. This unique RNA cluster 5 displays an inflammatory phenotype expressing many features of senescence and interferon signalling and response.

The mechanisms underlying senescence and IFN signalling are intertwined. IFN response can be triggered by a myriad of stimuli, including extra and intracellular double-stranded (ds) RNA and DNA from cell stress and apoptosis, cytosolic DNA, viruses, and microbes, which in turn activates a pro-inflammatory response.^93^ Over the course of aging, IFN pathways have been found to become aberrantly activated, which leads to global inflammation (*inflammaging*) and senescence.^94^ The presence of an IFN signature in brain cells is increasingly being associated with aging and neurodegenerative diseases in rodent models and humans.^95–98^ Type I IFN signatures are found upregulated in the aged brain, especially the choroid plexus, and in neurodegenerative diseases, where it may lead to recruitment and activation of immune cells and eventual neurodegeneration. Interestingly, in our model system, we identify hypomethylation of genes associated with IFN signalling in PMS fibroblasts, indicating a potential predisposition for developing an IFN response in people with PMS. Once fibroblasts are reprogrammed to iNSCs, they take on an even more pronounced IFN phenotype associated with senescence gene expression.

From the single-cell data, we establish not only the heterogeneity of iNSCs derived from both Ctrl and PMS donors, but also define a novel subset of disease associated RG-like cells that could be the ‘drivers’ of the inflammatory signature seen in the analysis of bulk PMS iNSCs. In fact, through ligand/receptor predicted interactions, the secretory factors from this – PMS mostly – IFN-responsive and senescent-like RNA cluster 5, could induce inflammatory-associated signatures (*i.e., neurodegeneration, oxidative stress induced senescence, and signalling by Ils*) and inhibit NSC maintenance (*i.e., Wnt signalling, and cell cycle*) in the other iNSC clusters, thus suggesting that RNA cluster 5 can further affect surrounding cells via amplification of such inflammation. Independent work from our group, further supports a key role for disease associated paracrine factors in conditioned iNSC media to induce neurite retraction and neuronal apoptosis.^46^

While the mechanisms of this RG-intrinsic intrinsic dysfunctional phenotype in PMS are still unknown, however, we do find epigenetic signatures starting even with the donor fibroblasts. Recently, neurodegenerative diseases have been found to be highly associated with viral exposure,^99^ and in the case of MS the Epstein Barr virus (EBV) increases risk of disease.^100^ Viral exposure combined with chronic inflammation in PMS may induce global epigenetic changes, affecting cells such as fibroblasts, found to be stressed in people with MS.^101^ Furthermore, the activation of human endogenous retroviruses (HERVs) in unique cell populations are also associated with chronic inflammatory neurodegenerative diseases^102^ and brain injuries.^103^ HERVs can be activated via inflammatory stimuli and induce an IFN response, similar to that of a viral infection.

Epigenetically, we identify sites with gains of accessibility are associated with IFN-response and signaling as well as cytokine production in RNA cluster 5 (corresponding to ATAC cluster 8) composed mainly of PMS derived cells. The epigenetic remodelling in this cluster was also associated with increased accessibility at binding sites of the p53 motif, known to be activated in response to cellular stress and DNA damage and promotes sustained IFN signaling and response.^85^

Using publicly available datasets from MS post-mortem brains, we assessed the expression of RG genes and identified non-neurogenic RG-like cells within well-known astrocyte clusters. Recent work has demonstrated that astrocytes exhibit plasticity in injury situations.^41^ Human pathologies which involved lesions and blood-brain barrier rupture were associated with a de-differentiation of astrocytes to replicating NSC/RG-like cells.^41^ Furthermore, this has been previously validated in rodent models where epithelial injury allows for neural precursors to dedifferentiate into multipotent NSCs in the olfactory epithelium.^104^ Based on these studies, astrocytes in PMS may be undergoing (i) a de-differentiation (or de-maturation) process, where they begin to express cell cycle and early RG-like cell markers due to exposure of chronic inflammation, and a (ii) resurgence as non-neurogenic RG-like cells at the level of disease-associated, ectopic, non-canonical niches.

Integration of our *in vitro* iNSC data with publicly available datasets in fact identifies and highlights a long-neglected, non-neurogenic disease-associated RG-like cell population, being found significantly increased in chronic active lesions areas and displaying IFN and senescence gene expression, which we term DARGs.

Interestingly, we identify twice as many DARGs in chronic active PMS lesions, which are slowly expanding in nature, feature smouldering inflammatory demyelination at the edge, remyelination failure, and axonal degeneration,^105^ and are associated with a more aggressive disease^106^. Further characterization of the phenotype of this novel DARG population showed increased expression of the SenMayo and IFN-associated genes compared to the astrocyte cluster.

Overall, our work shows there are epigenetic alterations in somatic fibroblasts isolated from people with PMS, and many of these epigenetic modifications remain following direct reprogramming into iNSCs. These epigenetic alterations are associated with de-repression (hypomethylation and increased chromatin accessibility) of IFN signalling and response as well as inflammation. We further uncover a novel subset of PMS iNSCs with high levels of inflammatory signalling, which we propose drives much of the bulk phenotype.

Lastly, we uncover a long-neglected DARG population in the PMS brain, which has similar transcriptomic profiles as the *in vitro* PMS iNSCs, including expression of senescence and IFN transcripts.

Our research lays the groundwork for further investigating ‘disease-pacemaker’ non-neurogenic RG-like cells in potentially driving neuroinflammation in neurodegenerative disease. Future work is needed to identify the origin and driver of epigenetic dysfunction arising in the cells of people with PMS.

## LIMITATIONS OF STUDY

While we generated cell lines from individuals with PMS, the variations in genetic backgrounds, sex, and age among these lines pose a potential limitation. Despite our thorough analysis of both patient and control cell lines, we acknowledge the necessity for additional validation of our findings in situ. Additionally, the inclusion of induced pluripotent stem cells (iPSCs) from the same donors would have enhanced the data quality, allowing for a more robust interrogation of the observed cellular phenomena. Single-cell spatial assays could offer a more comprehensive understanding of our results, particularly in capturing differences related to disease-relevant microenvironments and surrounding cells (neighbours).

## ACKNOWLEDGMENTS

The authors wish to acknowledge L. Bonfanti, A. D’Alessandro, V. Fossati, D. Franciotta, C. Frezza, O. Hruba, G. Pluchino, A. Quaegebeur, S. Rizzi, L. Roth and A. Speed for technical and intellectual inputs throughout this study.

This research was supported by the Ferblanc Foundation G112716 (SP and AMN); Catalyst Award from the UK MS Society H160 (SP and CMW); National MS Society Research Grant RG 1802-30200 (SP and LPJ); Bascule Charitable Trust RG98181 (SP); Wings for Life RG 82921 (SP and LPJ), and the Fondazione Italiana Sclerosi Multipla FIMS 2018/R/14 (SP and LPJ) and 2022/R-Single/011 (SP).

BP, DT, JW, LC, JL, JF, SD, MG, and IB are supported by the Intramural Research Program of the National Institute on Aging. AMN is the recipient of a European Committee for Treatment and Research in Multiple Sclerosis (ECTRIMS) Postdoctoral Research Fellowship Exchange Program fellowship (G104956) and is supported through a UK MS Society Centre Excellence grant (G118541). PP is supported through an MRC-DTP and Cambridge Trust PhD studentship and consumable award (RG86932) and Queen’s College Tutorial Award. RBI is supported through an MRC-DTP and Cambridge Trust studentship and consumable award (RG86932) and St. Edmund’s College Tutorial Award. CW is supported through a National MS Society Post-doctoral fellowship (FG-2008-36954). LPJ was supported by a Fondazione Italiana Sclerosi Multipla FIMS and Italian Multiple Sclerosis Association AISM Senior research fellowship financed or co financed with the ‘5 per mille’ public funding cod. 2017/B/5 (LPJ), a Wellcome Trust Clinical Research Career Development Fellowship (G105713), and a National MS Society Research Grant RFA-2203-39318. FE was supported by the Austrian Science Fund FWF (FWF-INTER – INTER/FWF/19/14117540/Pdage and the SFB F78 “Neuro Stem Modulation”, SFB F7810-B). LP and IM are supported by Wellcome Trust (203151/Z/16/Z) and the UKRI Medical Research Council (MC_PC_17230).

## AUTHOR CONTRIBUTIONS

Conceptualization, BP, AMN, DT, IM, SP, IB; Methodology, BP, AMN, DT, JW, LC, JL, RBI, JF, SD, MSC, AS, TL; Investigation, BP, AMN, DT, LP, PP, MLDN, LC, JL, RBI, CMW, GK, TL, MG, IM, SP, IB; Writing – Original Draft, BP, AMN, DT, LP, IM; Writing – Review & Editing, AMN, DT, IM, SP, IB; Funding Acquisition, AMN, CMW, LPJ, IM, SP, IB; Resources, JF, SD, MSC, AS, TL, FE; Supervision, MG, IM, SP, IB.

## METHODS

### Data and code availability

All codes used in the study are available on github: https://github.com/Core-Bioinformatics/DARG_PMS

- Any additional information required to reanalyse the data reported in this paper is available from the lead contacts upon request.
- All data generated for this study, in raw and processed format, are publicly available on the Gene Expression Omnibus (GEO), under accessions GSE243319, GSE251839, GSE251831, GSE251838, and GSE251830. Further data mining of processed data may be performed on bulkAnalyseR for bulk sequencing datasets and ClustAssess and Shiny Cell apps (https://bioinf.stemcells.cam.ac.uk/shiny/pluchino/DARG_PMS/bulkanalyser/, https://genomicspark.shinyapps.io/shinyApp/) for single-cell/ nuclei datasets. UCSC genome browser sessions for these datasets comprise: https://genome-euro.ucsc.edu/s/CSCI/DARG_PMS and http://tinyurl.com/2ygsa6or.

### EXPERIMENTAL MODEL AND SUBJECT DETAILS

#### Patient cells

The cohort consists of 5 PMS and 3 healthy controls between 25 and 63 years of age. The cohort includes representation from both genders, distributed across PMS and control groups (Table S1). PMS fibroblasts were provided by the New York Stem Cell Foundation (NYSCF) Research Institute through their Repository (http://www.nyscf.org/repository)^107^. Patients were recruited at the Tisch Multiple Sclerosis Research Center of New York, upon informed consent and institutional review board approval (BRANY). PMS donors underwent clinical assessment when recruited for the study. Control fibroblasts C1 and C2 (Table S1) were generated from adult dermal fibroblasts after obtaining consent and ethical clearance by the ethics committee of the University of Würzburg, Germany.

#### Generation and culturing of induced neural stem cells

iNSC lines were generated and quality controlled as described in *Ionescu, Nicaise et al.*^46^ and *Meyer et al.*^45^ iNSCs were maintained in neural induction media (NIM) consisting of DMEM/F12 and Neurobasal (1:1) (ThermoFisher), supplemented with N2 supplement (1x) (ThermoFisher), 1% glutamax (ThermoFisher), B27 supplement (1x) (ThermoFisher), CHIR99021 (3 µM) (Cell Guidance Systems), SB-431542 (2 µM) (Cayman Chemical), and hLIF (10 ng/ml) (PeproTech) until 70% confluent, then lifted using accutase, spun at 300 x g for 3 mins, and plated onto growth factor reduced (GFR) matrigel matrix coated plates (Corning) (1:20 in DMEM/F12) with Y-27632 (10 µM) (Miltenyi Biotec) between 1:3-1:5 in NIM media. Media was changed every second day as needed. Experiments were performed on cells from passages 20-35.

#### Fibroblast maintenance

Fibroblasts were maintained in growth medium (DMEM Glutamax I [Thermo Fisher]) supplemented with 10% fetal bovine serum, 1% non-essential amino acids and 1 mM sodium pyruvate (ThermoFisher) at 37°C with 5% CO_2_ and fed every 3-4 days. After reaching 90% confluency the fibroblasts were detached with trypsin-EDTA 0.05% for 5 min followed by neutralization in DMEM and spun down at 300xg for 5 min. They were split 1:4 into growth media onto tissue-culture treated plasticware.

### METHOD DETAILS

#### mRNA sequencing, analysis, and inference of Gene Regulatory Networks (GRNs)

iNSC lines, between passages 15-30, were plated at 500,000 cells per well in GFR-coated 6-well plates. After 24 hours, the media was refreshed with new NIM. Cells were harvested in RLT lysis buffer 72 hours after plating then frozen at −80°C until extraction. RNA extraction was performed according to standard steps described for the RNeasy kit, followed by DNase treatment (Qiagen). RNA was quantified using the NanoDrop 2000c instrument. Illumina Sequencing libraries were prepared using the TruSeq low sample protocol from 1 µg of total RNA (Illumina, San Diego, CA, USA). The resulting libraries were sequenced in paired-end mode, on 150 nts reads on an Illumina NovaSeq 6000 instrument.

The quality checking of the samples was assessed using fastQC v0.12.11, applied on raw files; the outputs were summarised using multiQC 1.14.^108^ Initial sequencing depths ranged from 30M to 44M reads; subsampling without replacement, done using seqtk 1.3-r106 ^109^, was performed to 34M reads, to avoid inconsistencies caused by uneven sequencing depths.^110^ All samples were aligned to the GRCh38.p13 genome using STAR 2.7.10a (paired-end mode).^111^ Expression quantification was performed using featureCounts v.1.6.3.^112^ The distribution of signal across transcripts was assessed on the UCSC genome browser. The tracks (bigwig format) were built from the bam files using samtools 1.17.^113^ Next, noisyR 1.0.0^48^ was used to estimate and remove noise from the count matrix; the raw expression levels were normalised using quantile normalisation.^114^ Differentially expressed genes (DEGs) were identified using edgeR^50^ and DESeq2^49^; due to the noise correction the DEG calling converged; p-values were adjusted using Benjamini-Hochberg multiple testing correction. bulkAnalyseR 1.1.0^47^ was used to build a shareable interface for the analysis and visualisation of data.

The Gene Regulatory Networks (GRNs) were predicted, and their dynamics assessed, on the bulk RNAseq data using a bulkAnalyseR ShinyApp.^47^ Additional analyses were performed using GENIE3^115^ and visNetwork^116^ to visualise subgraphs according to selected pathways and a maximum of 30 edges. To assess the global trend in co-variation of expression, for genes annotated to the selected pathways, density plots were created, per pathway, on the weights of the edges in the larger GRNs (corresponding values in the global adjacency matrix).

#### Immunoblotting

iNSCs were homogenized in 10X RIPA buffer (Abcam) supplemented with 100X protease and phosphatase inhibitors (ThermoFisher). Protein concentration was assessed using a BCA assay (ThermoFisher). Equal protein amounts (25 µg) were resolved by SDS-PAGE on Bolt™ Bis-Tris Plus pre-cast 4-12% gradient gels (Invitrogen) and transferred to 0.45 mM polyvinylidene fluoride (PVDF) membranes (Thermo Scientific). Membranes were blocked with TBS blocking buffer (LI-COR Biosciences) and immunoblotted with the indicated antibodies: mouse anti-p16^Ink4a^ (Invitrogen) at 1:500, rabbit anti-GDF15 (Proteintech) at 1:1000, and mouse anti-b-actin (Sigma) at 1:5000 in TBS blocking buffer (LI-COR Biosciences) with 0.1% Tween, followed by fluorescent secondary antibodies IRDye 680RD Goat anti-Rabbit or IRDye 800CW Goat anti-Mouse (LI-COR Biosciences) at 1:10,000 in TBS blocking buffer (LI-COR Biosciences) with 0.1% Tween and 0.01% SDS. The immunoblots were visualized with the ChemiDoc MP Imaging system (Bio-Rad). Densitometric analysis was conducted with Fiji by ImageJ. Protein targets were normalized to β-actin.

#### SPiDER-gal

Cells were plated on black-walled, clear bottom 96-well plates (ThermoFisher, 165305) at 15,000 cells/well and maintained in culture for 5 days. Expression of senescence associated β-galactosidase was measured by a SPiDER-β-gal-based cellular senescence plate assay kit (Dojindo) according to manufacturer’s instructions. Briefly, cells were washed with PBS, stained with 1 μg/mL Hoechst (Sigma-Aldrich) as a measure of cell number, washed again, before the fluorescence intensity was measured at 358nmEx/461nmEm. Cells were then lysed with the provided buffer and the SPiDER-β-gal stain was added and incubated at 37°C overnight. Fluorescence intensity was measured at 520ex/565em. The SPiDER-β-gal fluorescence intensity of each well was corrected for the autofluorescence of empty wells and normalized to the Hoechst fluorescence intensity of the respective well to normalize for cell number. The resulting average SPiDER-β-gal/Hoechst fluorescence intensity of each cell line was normalized to that of healthy control cell lines.

#### Cell Cycle Analysis

iNSCs were plated at a density of 80,000 cells/cm^2^ on GFR-coated plates. After 3 days cells were lifted using accutase and then pelleted at 500 x g for 5 minutes. The cells were fixed in 70% ethanol for 30 minutes on ice then pelleted at 850 x g for 5 minutes. The cell pellet was resuspended in RNase (100 ug/mL) for 15 minutes and incubated at room temperature. Propidium iodide (1 ug/mL) was added to each sample and cells were analysed on a BD LSRFortessa with the flow rate on slow. 20,000 events were collected for each sample. The data was analysed using FlowJo 10.9 software using the Dean-Jett-Fox approach.

#### Telomere length analysis

Relative telomere length was assessed using the *Joglekar et al*. protocol using quantitative PCR (qPCR) and comparison to that of a single copy gene.^117^ iNSCs were plated at a density of 80,000 cells/cm^2^ on GFR-coated plates. After 3 days cells were lifted using accutase and then pelleted at 500 x g for 5 minutes. DNA was isolated according to the DNeasy Blood & Tissue Kit (Qiagen) and quantified using the Nanodrop 2000c instrument. For initial optimization of the qPCR reaction, the DNA was diluted to three different concentrations (100 ng/μL, 25 ng/μL, 6.25 ng/μL), and it was determined that 100 ng/uL had the best efficiency for both the human β-globulin and telomere primers. Two PCR reactions were separately conducted, for human β-globulin the mastermix was made using 5 μL Fast SYBR Green (ThermoFisher), 1 μL of hbg1 primer (3 μM), 1 μL of hbg2 primer (7 μM), and 2 μL of nuclease-free water. The reaction was cycled at 58°C annealing temperature along with a melt curve analysis using a QuantStudio 7 Flex (ThermoFisher). For the telomere primers, the mastermix was made using 5 μL Fast SYBR Green (ThermoFisher), 1 μL of telomere A primer (1 μM), 1 μL of telomere B primer (3 μM), and 2 μL of nuclease-free water. The reaction was cycled at 56°C annealing temperature along with a melt curve analysis using a QuantStudio 7 Flex (ThermoFisher). Each sample was run in duplicate. Average telomere length was calculated as the ΔΔCT = (PMS average hbg Ct – PMS average telomere Ct) – (Control hbg Ct – control average telomere Ct).

#### Whole genome bisulfite sequencing (WGBS)

Genomic DNA was extracted from 100,000 fibroblasts and iNSCs using the DNeasy Blood and Tissue Kit (Qiagen). The quantity of DNA was measured using the Quant-iT PicoGreen method and victor X2 fluorometry (ThermoFisher), and the integrity of the DNA was evaluated with Agilent genomic DNA screen tape. 500 ng of genomic DNA was used for sequencing. The sample quality control criteria for the WGBS library were set to having a DNA integrity number (DIN) score of 7.0 and above. The extracted DNA was fragmented to an average insertion size of 550 base-pairs and the fragments were attached to end-repaired adapters. Genomic DNA was bisulfite converted using the EZ DNA methylation Gold kit (Zymo, Catalog #D5005) following the manufacturer’s instructions. We next applied the xGen Methyl-Seq Lib Prep kit (Integrated DNA technologies, Catalog #10009824) to the prepare the genomic DNA library. Library quality control was performed using qPCR (LightCycler 480) and TapeStation 4200 (D1000 screen tape).

The dataset comprises 8 samples (4 fibroblast lines and 4 iNSC lines), with sequencing depths varying from 213M to 408M, and an average of 315M reads per sample. Reads with adapter contamination were trimmed using Trim Galore (0.4.3)^118^ with options: --paired –q 25. Trimmed reads were aligned to the *H sapiens* reference genome (version hg38), using HISAT2^119^ (version built in the current stable version of Bismark 0.23.1^120^). A bisulfite-converted index (GA and CT conversion) was generated with default parameters. We identified 28M CpG sites per sample, with sequencing coverage varying from 24x to 39x, (an average of 30x coverage per sample). The bismark_methylation_extractor tool was used to summarize the methylation levels at CpG sites. After assessing the bias at 5’end regions using M-bias results, the first 2nts were excluded, as follows: bismark_methylation_extractor -p –ignore 2 –ignore_r2 – comprehensive –no overlap –bedGraph –counts –buffer_size 16G ($Aligned read bam file).

#### Identification of differentially methylated sites and regions

MethylKit^121^ was used for DMC and DMR quantifications, and fibroblast *vs* iNSC comparisons. A minimum threshold minimum of 10nts coverage for downstream DNA methylation analysis was set. The aligned reads were split into 100nt tiles (DMRs) using metilene.^122^ Differential methylation was calculated, applying a McCullagh and Nelder^123^ correction for overdispersion, as well as the sliding linear model (SLIM) proposed in methylKit to correct for multiple testing. Tiles with a q-value < 0.05 and over 20% methylation difference were called differentially methylated. Motif enrichment analysis was performed using Homer (*findMotifsGenome.pl*). Annotations relevant for the hg38 v6.4 of the *H sapiens* reference genome (genes, exons, introns, UTRs, and other annotations) were extracted using Homer annotation tools. AnnotatePeaks.pl DMR hg38^124^ was used to evaluate the distribution of methylation across the genome. Next, a comparative analysis of the DMRs/DMCs across tissue types, contrasting the control and PMS samples was performed. In addition to the number of methylated tiles per annotation category was calculated, as well as their distance to the closest Transcription Start Site (TSS). To calculate the epigenetic age, we applied the Shireby-Cortex,^56^ Hovarth,^57^ and Zhang,^58^ ageing clocks frameworks. For the Hovarth and Zhang estimations Clockbase platform^125^ was used, relying on matched Illumina methyl array IDs. The DNA methylation levels (0-100%) per CpG probe, and the sample metadata were submitted to Clockbase, and the predicted clock age was downloaded as CSV format. For the Shireby-Cortex^56^ aging estimate, we downloaded the DNA methylation probes, and coefficient values and relied on matched Illumina methyl array IDs. We used total 347 DNA methylation CpG probes to predict epigenetic aging (> 20nts coverage).

#### Nuclei isolation, library preparation, and RNA sequencing

For the single-nucleus single-omics, (ATAC) and multiomics experiments, respectively, 2 ×10^6^ cells were harvested; nuclei were isolated following the manufacturer’s instructions with minor modifications. Briefly, cells were lysed in 100 μL of freshly prepared lysis buffer (1 mM Tris-HCl [pH 7.4], 1 mM NaCl, 300 μM MgCl_2_, 0.01% Tween-20, 0.01% IGEPAL CA-630, 0.001% Digitonin, 0.1% BSA, 100 μM DTT, and 100 mU/μL RNase inhibitor) for 1 minute on ice, washed twice in 500 μL of wash buffer (1 mM Tris-HCl [pH 7.4], 1 mM NaCl, 300 μM MgCl_2_, 0.01% Tween-20, 0.1% BSA, 100 μM DTT, and 100 mU/μL RNase inhibitor), and the number of nuclei was assessed using the Countess II FL Automated Cell Counter (ThermoFisher). Thereafter, approximately 16,000 nuclei were incubated with the transposase enzyme, loaded into Chromium Next GEM Chip H Single Cell Kit (10x Genomics). snATAC libraries were generated using Chromium Single Cell ATAC Reagent Kits User Guide v1.1 (10x Genomics) according to manufacturer’s instructions; for the multi-omics samples, nuclei were loaded into Chromium Next GEM Chip J Single Cell Kit (10x Genomics); libraries were prepared using Chromium Next GEM Single Cell Multiome ATAC + Gene Expression Reagent Kits (10x Genomics) according to manufacturer’s instructions. The quality of the libraries was checked on the Agilent Bioanalyzer with High Sensitivity DNA kit (Agilent); per sample libraries were sequenced on Illumina Novaseq 6000 with target sequencing depths of 25,000 -70,000 reads per nucleus.

For single-cell (sc)RNAseq, cells were counted using a hemocytometer, 10,000 cells were loaded into Chromium Next GEM Chip G Single Cell Kit (10x Genomics), and scRNA libraries were generated with Chromium Single Cell 3’ Reagent Kits v3.1 (10x Genomics) according to manufacturer’s instructions. The quality of the libraries was checked on the 4200 Agilent Tapestation with High Sensitivity DNA kit (Agilent); per sample libraries were sequenced on Illumina Novaseq 6000 with target sequencing depths of 30,000 - 65,000 reads per cell.

#### Single-cell transcriptomics/ epigenetics data pre-processing

Standard CellRanger pipeline (6.1.2) and CellRanger ARC (2.0.0) were applied for aligning reads to the aforementioned version of the *H sapiens* genome and for quantifying gene/ peak expression. For the RNA component, intron-matching reads contributed to the gene expression levels. The processed gene expression matrix was imported in Seurat.^126^ Additional filtering was performed on the distributions summarizing the number of counts, features, and percentages of reads incident to mitochondrial and ribosomal genes, across cells, per sample (accepted cells satisfied the criteria: number of UMIs > 8,000, number of genes per cell > 1,000, log_10_ (genes per UMI) > 0.75). We observed different ranges for MT and RP proportions for the scRNAseq and snRNAseq samples, respectively (we retained cells with 15-40% RP [scRNAseq] and 2-25% RP [snRNAseq]). Post-filtering, on all retained cells, the MT and RP entries were excluded from the expression matrix, pre-normalization. The normalization of expression levels was based on log_2_ normalization (scale.factor = 10000). The cell cycle assignation was performed in Seurat using the ‘CellCycleScoring’ function and *a priori* defined gene set.

#### Clustering

Next, we applied the ClustAssess framework to determine optimal, data-driven parameters, starting with the number and type features according the stability of resulting partitions.^70^ We used Element-centric similarity,^70^ summarized on 30 iterations into Element centric consistency (ECC),^71^ to objectively assess stability.^126^ Highly variable features (N=1,000 determined using the vst) approach, yielded optimal outputs. A 20-shared nearest neighbour (SNN) graph was constructed on the HGV PCA.^126^ To address batch effect across sc and sn quantifications, we applied Harmony.^127^ Clustering was performed using Louvain approach (resolution=0.2), implemented within Seurat^126^ v4.0.5. 8 clusters produced a stable partition on scRNAseq and snRNAseq components. We excluded the smallest cluster (133 driven by a specific sample (i.e. C1-specific cluster). A X^2^ test was used to assess the significance of proportions of cells for PMS *vs* control samples. Marker genes (on cluster *vs* complement and pairwise differential expression) were identified using the ‘findMarker’ function. The top 5 most positively differentially expressed genes were visualized in a heatmap.

Enrichment analyses were performed using gprofiler^128^ on markers called on a Wilcox test with a |log_2_(FC)| threshold of 0.25, an adjusted p-value (Benjamini Hochberg multiple testing correction) less than 0.05 and a minimum percentage of cells expressing the gene of 0.1, in either subset. The background set for the enrichment analysis comprised all genes expressed in at least 10 cells.

#### Voting Schemes

A variable voting-scheme was used to identify cell subsets requiring a minimum number of expressed genes corroborated with a minimum average expression level. The voting scheme for radial glial gene signature includes cells expressing 6 out of 9 manually curated genes (**Table S4**) with a normalized expression threshold of 1; glial progenitor cells express 7 out of 10 genes (**Table S4**) with an expression threshold of 0.5; neural progenitor cells express 5 out of 7 genes (**Table S4**) with an expression threshold of 0.5. The voting scheme for IFNα/β signalling is based on cells expressing 3 out of 6 genes (**Table S4**) with an expression threshold of 0.5; NOTCH1 signalling comprises cells expressing 13 out of 16 genes (**Table S4**) with no abundance threshold (I.e. presence/ absence).

#### Re-analysis of *Absinta et al.* and *Schirmer et al.* Data ex vivo datasets

Raw fastQ files from the studies by *Schirmer et al.* (2019)^77^ and *Absinta et al.* (2021)^16^ were downloaded from ENA using fasterq-dump. The quality checking and mapping leading to the filtered feature-barcode matrices were performed as described above. For the *Schirmer et al.* (2019) dataset, cells with less than 4,000 features/genes were retained; an upper bound of 15,000 was employed for the maximum number of UMIs per cell; 5% is maximum proportion of fragments incident to mitochondrial DNA; 10% is the maximum proportion of reads incident to nuclear ribosomal genes. For *Absinta et al.* (2021), cells with the number of features between 200 and 5000 were kept for subsequent steps of the analysis; an upper threshold of 20,000 UMI counts was used, and the maximum mt% was set to 5%. Both datasets were normalized using SCTransform;^129^ the *Absinta et al.* (2021) dataset was batch corrected using Harmony^127^ on the patient variable, with θ = 2. To detect stable partitions, on each separate dataset, ClustAssess^70^ was used with 20-50 iterations, assessing resolution parameters between 0.1 and 1.5 (0.1 increment steps). For the *Schirmer et al.* (2019) dataset, the top 4500 highly variable features yielded the most stable partitions; for the *Absinta et al.* (2021) dataset the top 3,500 highly variable features were selected. For both, the optimal resolution value was 0.6.

#### Pseudotime Analysis

Monocle3^130^ was used to infer trajectories for the *in vitro* data, as well as for the single-cell data from the *Schirmer et al.* and *Absinta et al*. studies. To identify the start and the endpoint, a selection of genes was used in a voting approach. The manually curated set of genes, used for determining the starting region in the *in vitro* data, comprises TOP2A, CENPF, UBE2C, ASPM, APOLD1 with an expression threshold of 2 and a tolerance of 1 gene, i.e. any one gene from the set maybe not expressed; for the ending region, genes IFIT2 and CDKN2A were used, with an expression threshold of 1.5 and a tolerance of 0.

For the *Absinta et al.* and *Schirmer et al.*, larger subsets of genes were used; for the former, the following genes were used for the starting region: *LY6E*, *PPAN*, *FASN*, *CLU, SORD*, *TRAP1*, *TUBB2A*, *AP1S2*, *YBX3*, with an expression threshold of 0.5 and a tolerance of 4 genes; for the ending region, the following genes were used *ISG15*, *B4GALT5*, *IFITM3*, *SAR1A*, *KIAA1217*, *TRPC4*, *FGF4*, *B2M*, *ZC3HAV1*, *WARS*, *FN1*, *IFIT1*, with an expression threshold of 0.5 and a tolerance of 4 genes missing. For the *Schirmer et al* dataset, the ending region was defined by genes *ISG15*, *B4GALT5*, *IFITM3*, *SAR1A*, *KIAA1217*, *TRPC4*, *FGF4*, *B2M*, *ZC3HAV1*, *WARS*, *DDX58*, with an expression threshold of 0.5 and a tolerance of 6 genes.

Using the ordering of cells based on their transcriptomic signatures, the ClustAssess stability framework was applied on gene expression levels. This yielded a stable number of gene-clusters, named gene modules, representing a precursor of GRN inference; the genes per module were further characterized from a pathway perspective (using gprofiler^128^), against GO terms, KEGG and REAC terms, and functional elements (TFs and miRNAs). Next, we chose three gene modules that characterized sections of interest on the trajectory-based UMAPs of the *in vitro* dataset. The genes within each module were used to create a proxy (a transcriptomic pattern) subsequently employed to identify homologue gene modules computed based on the *ex vivo* datasets, *Schirmer et al.* and *Absinta et al.,* respectively. Briefly, we considered the percentage of genes present in the *ex vivo* gene modules using the three *in vitro* gene modules; to account for the variable number of genes for bo*th in vitro* and *ex vivo* modules the outputs are scaled by the size of the gene set, i.e. larger gene sets are penalized more than smaller gene sets. The pairwise comparison of gene modules (**Fig. S4A-D**) relies on Fisher’s exact tests, using the *in vitro* data as baseline comparator. Benjamini-Hochber (FDR) correction was applied to account for the multiple testing pyscenic^131^ was used to infer regulatory interactions, aligned with the metadata available for the *Homo sapiens* (hg38) reference genome. A docker container was used to generate a loom object from the existing Seurat object. Loompy was used^132^ to create a SCope object, explored using the SCope web application; figure 6G illustrates specific regulons.

#### Cell-cell regulatory interactions and effects

NicheNet^80^ (v 2.0.4) was used to predict intercellular regulatory interactions, based on ligand-receptor databases (weighted_networks_nsga2r_final.rds). The correlative analysis, summarized as interaction scores, was applied on cluster-specific marker genes (differentially expressed genes). Further analyses were focused on cluster 5 (“inflammatory cluster”) assigned as sender cells *vs* receiver cells, as the remaining clusters, respectively. The analysis was performed separately on control and PMS iNSCs, respectively. The summary of interactions was visualized using circos plot (circlize library v.0.4.15).

#### Cytokine Array

iNSCs were plated at a density of 100,000 cells/cm^2^ on GFR-coated plates. Media was collected on day 5 from each line. The Human Cytokine Antibody Array C5 (RayBiotech) was used for semi quantitative detection of 80 proteins according to manufacturer’s instructions.

Overnight incubation was performed for steps when the option was given. Membranes were exposed using a Gel Doc XR imager (BioRad). Blots were analyzed using the Protein Array Analyzer macro for ImageJ (written by Gilles Carpentier, 2008). The relative quantity of each protein was normalized to the positive and negative controls included on the array. The array was performed once for each iNSC line. Control lines were averaged together to generate a fold change comparison over PMS iNSC lines. To visualize the results, we calculate the Z-score per cytokine using the heatmap function in an R environment.

#### snATACseq analysis

CellRanger ARC2.0.0 (multi-omics) and CellRanger ATAC2.0.0 (snATAC only) were used to map reads and quantify expression for the single nuclei ATAC-seq datasets. The peak calling was performed on pseudobulked input, comprising cells with at least 100 reads sequencing depth. Union peaks (peaks present in at least one sample) were reported. We excluded peaks overlapping the ENCODE-defined blacklist regions (hg38). To address the variation in sequencing depths, across samples, we normalized expression levels using random subsampling without replacement.^110^ The set of fragments (with lengths varying from 200 to 400 nts) *vs* the union-peaks were used to generate the ATAC expression matrix. For downstream analysis we relied onSeurat^126^, Signac^133^, and ArchR^134^ packages. Additional quality controls include assessment of nucleosome signatures and TSS enrichment analysis. we filtered the fragments with nucleosome signals < 4 and TSS enrichment levels > 2. Peak intensities were normalized using the term frequency inverse document frequency (TF-IDF) normalization (scale factor = 10,000). The dimensionality reduction was performed using latent semantic indexing (LSI). Additionally, we performed Harmony integration across batches, which was used as input for the final clustering (resolution=0.2, SLM method^126^). Differentially expressed peaks were identified using the ‘findMarker’ function (Seurat package). We performed *de novo* motif analysis using Homer (*findMotifsGenome.pl*) and GO term enrichment analysis using GREAT with the background of the whole genome.^135^

#### Integrative analysis of multimaps profiles

The integration of snRNAseq and snATACseq signals was performed on 5,242 cells with matched barcodes. The crosstalk between modalities was assessed using the partitioning information obtained on single modalities. The co-variation in expressed was summarized in joint ATAC/RNA heatmaps, with Z scores, calculated per modality, on pseudobulked expression per gene being presented for the gene itself (RNA modality), TSS proximal peaks (<3kb) and TSS distal peaks (greater than 3kb and less than 50kb). Both ATAC and RNA modalities were used to infer regulons using SCENIC+.^136^

#### Statistical analysis

For all phenotypic analyses, a p-value < 0.05 was considered significant (*). We performed statistical tests described in individual figure legends using Prism software version 10 (GraphPad Software, San Diego CA). A Benjamini-Hochberg, False discovery rate (FDR) multiple testing correction was applied to account for Type I errors. For low throughput differential expression analysis on genes, we used a negative binomial test with the FDR cutoff value set to <0.05.

All analyses were performed on R 4.2.3, on high memory computer (MacPro M1 Max, 64GB memory) and servers (Intel E7-8860v4, 3TB memory).

