## Supplemental Figures for "Integrative single-cell analysis of neural stem/progenitor cells reveals epigenetically dysregulated interferon response in progressive multiple sclerosis"

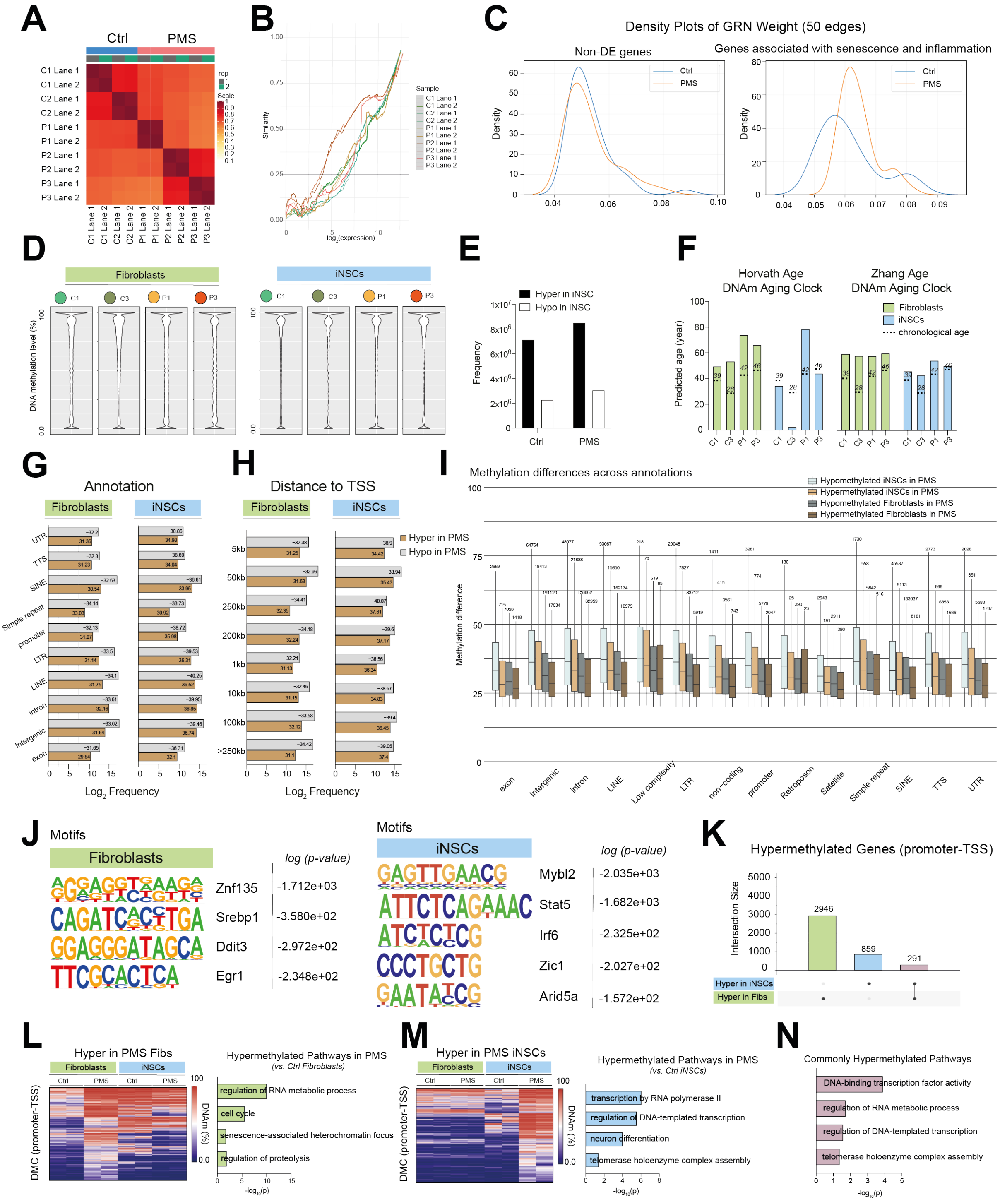


### Figure S1. Supporting analyses on mRNAseq and WGBSseq

(**A**) Jaccard similarity index summarizing the experimental design of the bulk RNAseq datasets.

(**B**) Identification of noise threshold using the noisyR framework.

(**C**) Density plots of the covariance in gene expression (derived from gene regulatory network (GRN) inference using GENIE3); networks inferred on Ctrl (blue) and PMS (red) samples, with top 50 edges considered were built on non-differentially expressed genes (left) and differentially expressed genes associated with senescence and inflammation (right).

(**D**) Summary of DNA methylation levels on fibroblast and iNSC samples.

(**E**) Frequency of differentially methylated cytosines, (hyper and hypomethylated), in Ctrl and PMS cells, plotted as fibroblasts *vs* iNSCs.

(**F**) Horvath and Zhang predicted DNA methylation aging clock results on parental fibroblasts and directly reprogrammed iNSCs.

(**G**) Frequency of hyper and hypomethylation of PMS fibroblasts and iNSCs *vs* Ctrl based against gene annotation classes.

(**H**) Frequency of hyper and hypomethylation of PMS fibroblasts and iNSCs *vs* Ctrl based on distance to TSS.

(**I**) Hyper and hypomethylation differences across annotations classes for PMS fibroblasts and iNSCs *vs* Ctrl. Numbers represent identified DMRs.

(**J**) HOMER *de novo* motif enrichment analysis of PMS fibroblasts and iNSCs from hypomethylated regions in (**Fig. 2D**) representing unique motifs to PMS fibroblasts and iNSCs. TF, transcription factor. The motifs are represented using proportional sequence logos.

(**K**) UpSet plot of hypermethylated genes in fibroblasts and iNSCs within the 3k upstream of the transcription start site (TSS) - defined in this context as the promoter region.

(**L-M**) Heatmap and enrichment analysis of downstream genes of hypermethylated DMCs specific to PMS fibroblasts *vs* Ctrl and PMS iNSCs *vs* Ctrl.

(**N**) Enrichment analysis of commonly hypermethylated genes of PMS fibroblasts and iNSCs *vs* Ctrl.

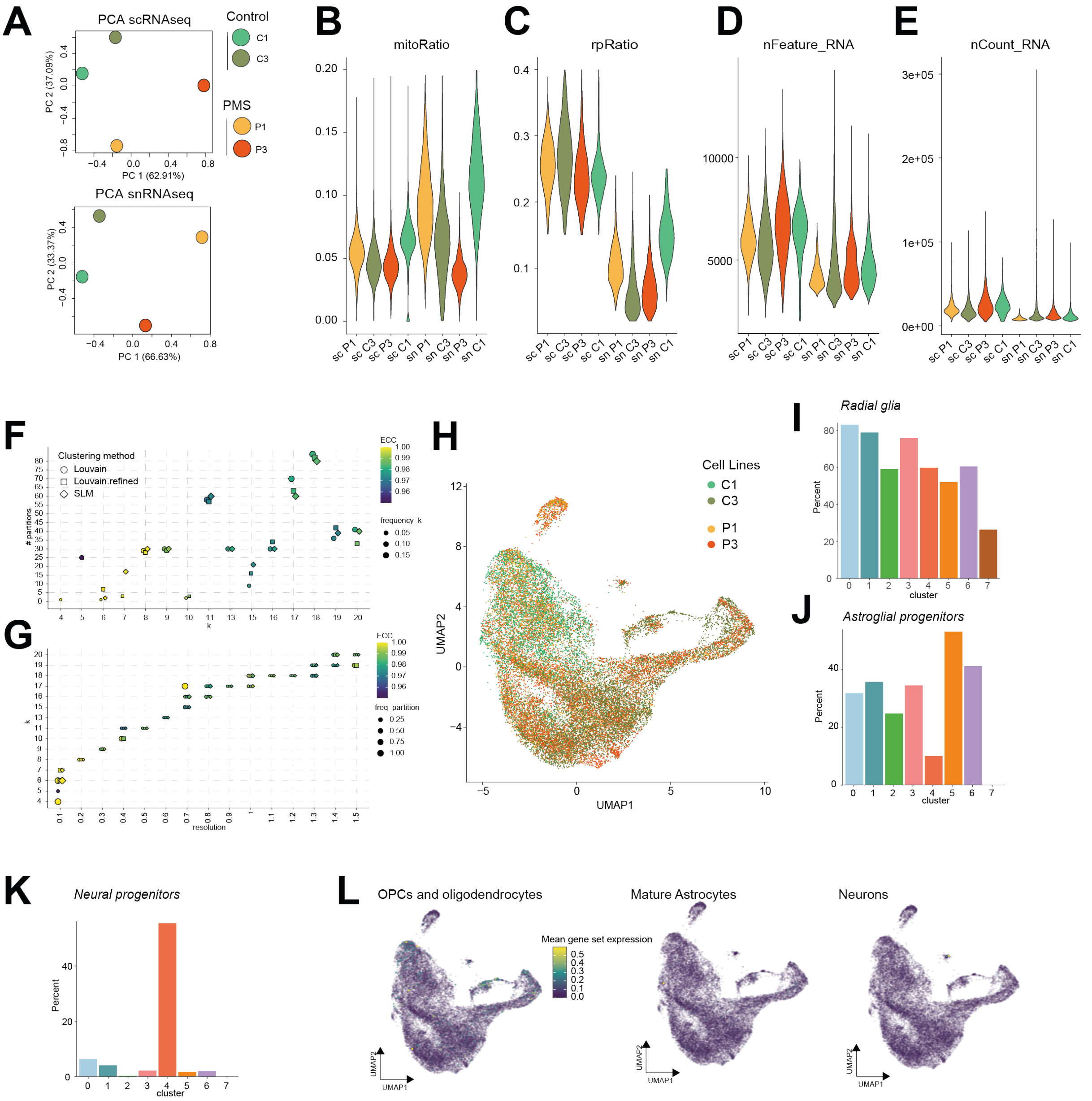


### Figure S2. Quality control of sc/ snRNA sequencing dataset

(**A**) PCA plots summarizing pseudobulked co-variation in expression of samples from iNSCs.

(**B**) Distribution of proportions of reads incident to mitochondrial genes per sample. A threshold of 20% was applied for filtering putative apoptotic cells.

(**C**) Distribution of proportion of reads incident to ribosomal proteins across samples; a threshold of 40% was applied; we also note a striking difference between single-cell and single-nuclei samples. MT and RP genes were excluded from downstream analyses prior to normalization.

(**D**) Distribution of nFeature (the number of genes per cell) in sc and snRNAseq dataset

(**E**) Distribution of nCount (the number of transcripts per cell) in sc and snRNAseq dataset

(**F**) ClustAssess summary figure illustrating the clustering stability, linking k (number of partitions correspondence with ECS threshold = 1, x-axis) *vs* the number of partitions (y-axis) derived from three clustering algorithms.

(**G**) ClustAssess summary figure illustrating cluster stability *vs* resolution (x-axis) and the number of clusters (y-axis).

(**H**) RNA UMAP illustrating the distribution of cells across the individual samples.

(**I-J**) Histograms representing percent of cells from individual cluster based on cell type, radial glia (**I**), astroglial progenitors (**J**), and neural progenitors (**K**).

(**L**) UMAPs of mean gene set expression for oligodendrocyte progenitor cells (OPCs) and oligodendrocytes, mature astrocytes, and neurons in the iNSCs.

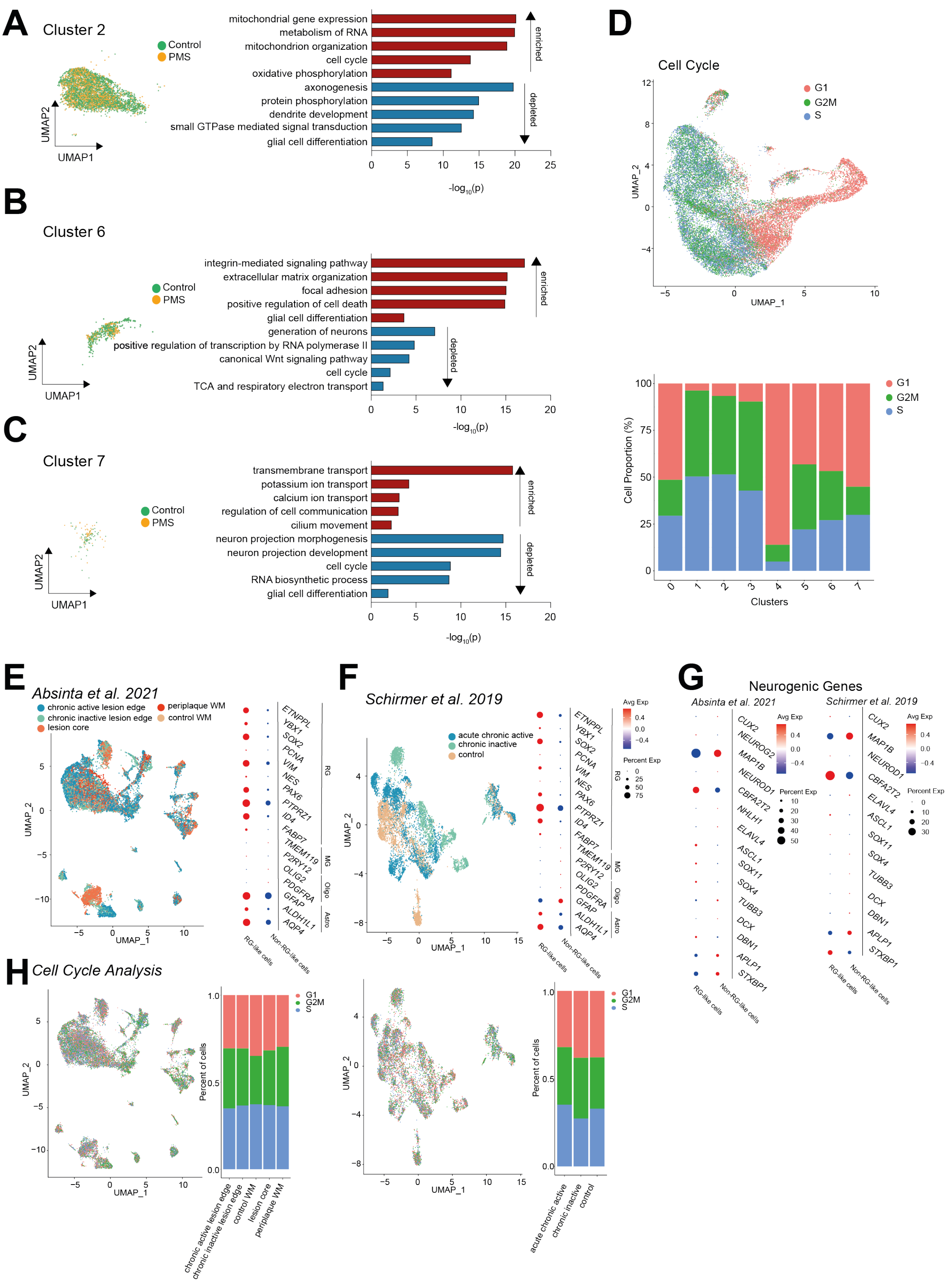


**Figure S3. iNSCs do not contain differentiated cells and RG-like cells in the adult human brain are not neurogenic**

**(A-C)** Enrichment analysis depicting enriched and depleted terms in cluster 2; cluster 6; cluster 7 *vs* all other clusters.

(**D**) UMAP summarizing cell cycle allocation (G1, G2M, and G) and stacked histogram summarizing proportions of cells per cluster assigned to each cell cycle stage.
(**E**, **F**) Recalculated UMAP based on RG gene expression displaying cells based on lesion location. Bubble plots displaying average expression and percent expression of RG markers, microglia (MG) markers, oligodendrocyte markers, and astrocyte markers in the identified DARGs and non-RG-like cells (**Fig. 3L-M**). WM, white matter.
(**G**) Bubble plots of neurogenic genes in RG-like and non-RG-like cells.
(**H**) UMAP summarizing cell cycle allocation and stacked histogram summarizing proportions of cells per area sequenced to each cell cycle state in the *ex vivo* data.

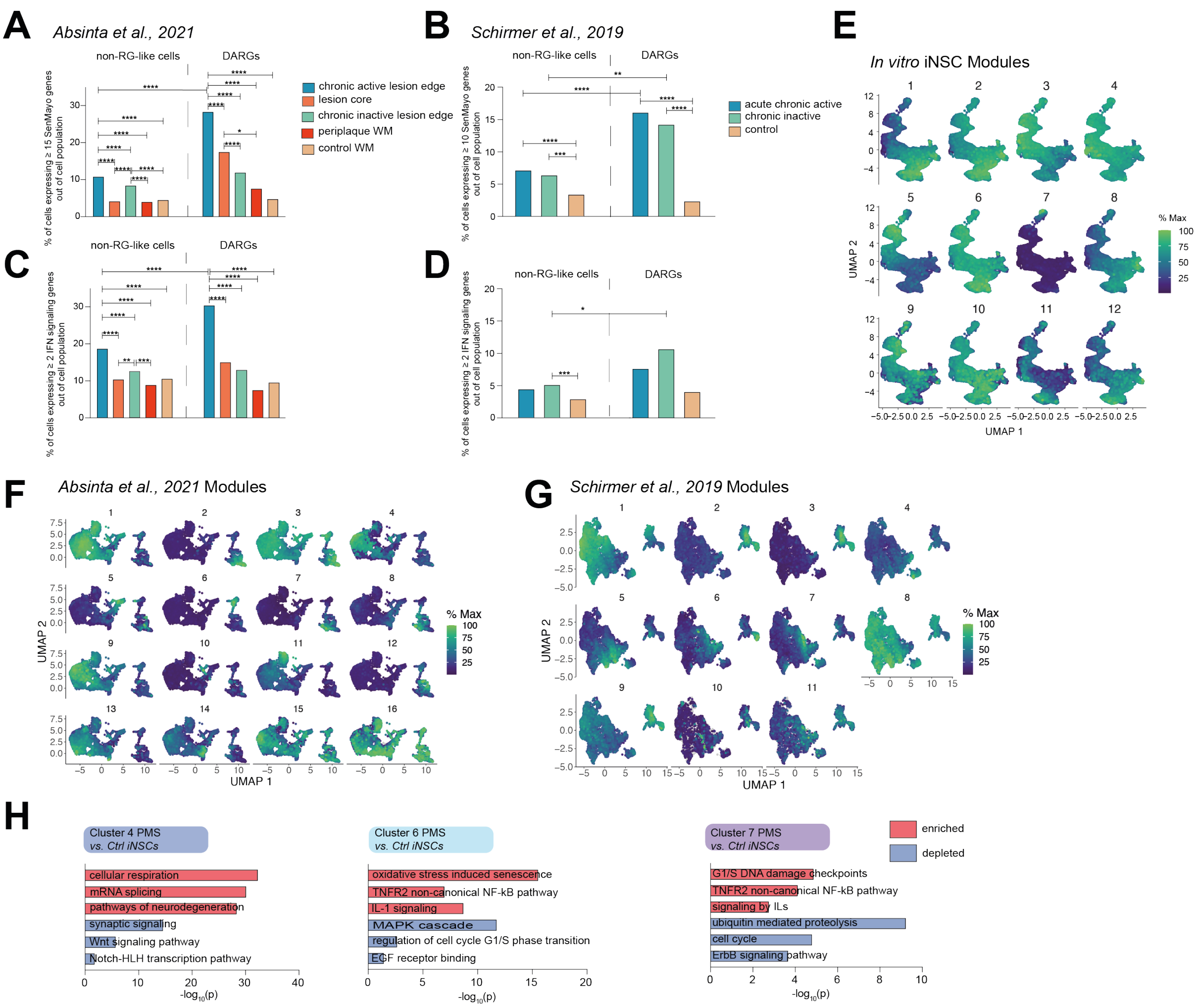


**Figure S4. DARGs in chronic active lesion edges display senescence and IFN signaling genes**

(**A**) Relative proportion of cells expressing ≥ 15 SenMayo genes for DARGs and non-RG-like cells in the *Absinta et al*., *2021* dataset, summarized per brain area sequenced.

(B) Relative proportion of cells expressing ≥ 10 SenMayo genes for DARGs and non-RG-like cells in the *Schirmer et al., 2019* dataset based on brain area sequenced.

(C-D) Relative proportion of cells expressing ≥ 2 IFN signaling genes for DARGs and non-RG-like cells in the *Absinta et al., 2021* (C) and *Schirmer at al.*, *2019* (D) dataset based on brain area sequenced.

(E) Gene modules generated on (dataset specific) inferred pseudotime from iNSC *in vitro* data.

(F-G) Gene modules generated on inferred pseudotime form *Absinta et al., 2021* and *Schirmer et al., 2019* recalculated UMAPs.

(H) Functional summary of enriched and depleted genes in cluster 4, cluster 6, and cluster 7 (Ctrl *vs* PMS, *in vitro* data).

Data in A-D are represented as percentages. Adjusted-p values, with a Benjamini-Hochberg multiple testing correction: *p≤ 0.05, **p≤ 0.01, ***p≤ 0.001, ****p≤ 0.0001 resulting from a Fisher's exact test on all pairwise comparisons.

>=

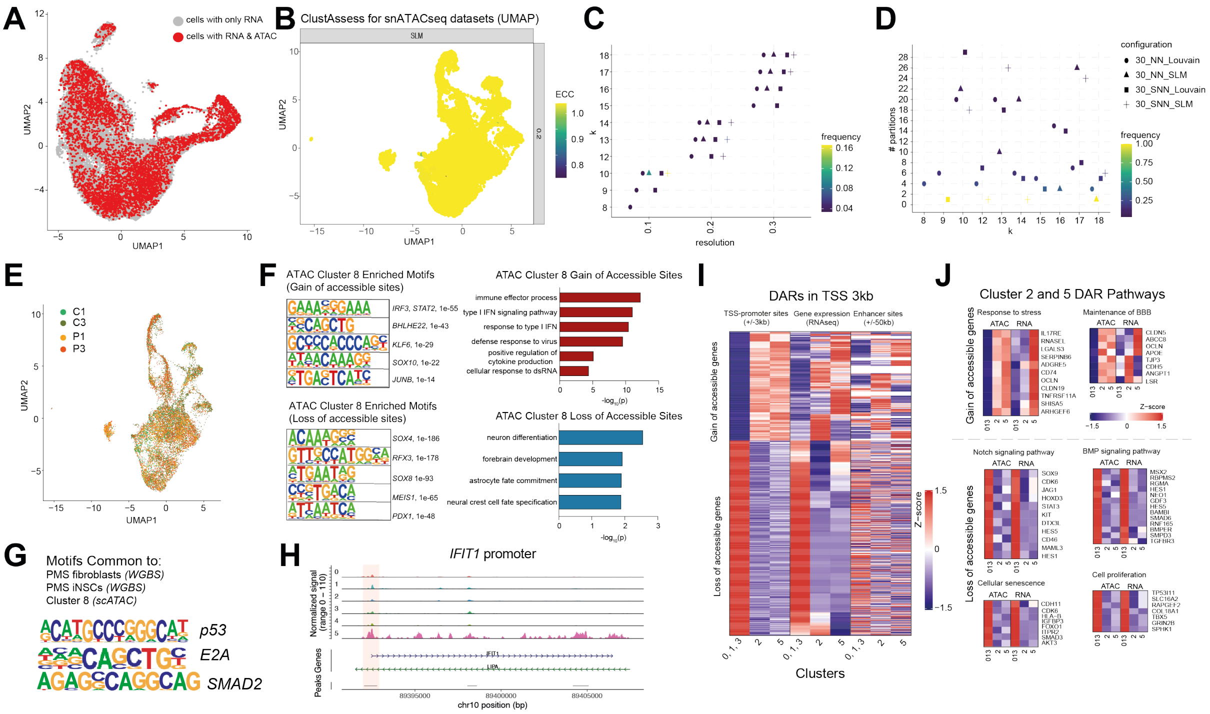


### Figure S5. Summary of changes in chromatin accessibility specific to the inflammatory cluster. Assessment of functional variance on regulatory regions

(**A**) UMAP of matched cells (red) with both RNA and ATAC expression and accessibility signatures, resulting from the multi-omics dataset.

(**B**) ClustAssess summary of ECC, represented on the ATAC UMAP, indicating a high stability of ATAC-centric clusters.

(**C-D**) Identification of stable, data-driven configurations for ATAC clusters, on 30 nearest neighbours (on weighted and unweighted neighbourhoods) for variable resolution (**C**) and number of clusters (**D**).

(**E**) ATAC-centric UMAP summarizing the uniform distribution of cells, across all samples.

(**F**) *De novo* motif analysis from gain of accessibility (GA) and loss of accessibility (LA) regions in ATAC cluster 8, summarized as proportional sequence logos; GREAT prediction of cis-regulatory regions using GA and LA regions in ATAC cluster 8.

(**G**) Summary of motifs, represented as proportional sequence logos, enriched in PMS fibroblasts and iNSCs on the WGBS modality, and in cluster 8 via scATACseq.

(**H**) Genome browser tracks illustrating gain-of-accessible sites within *IFIT1* in RNA cluster 5.

(**I**) Heatmap summarizing differentially accessible regions (DARs) in 3kb regions upstream of the TSS. Heatmap includes the promoter region, the coding region, and enhancer sites for genes part of RNA-clusters 0,1,3, cluster 2, and cluster 5. The genes are sorted on loss/ gain of accessibility, and on expression patterns.

(**J**) Dual heatmaps structured based on enrichment analyses, and allocation to pathways, summarising the variation in expression and accessibility for genes with gain and loss of accessible sites, in RNA-clusters 2 and 5.
